# CALDERA: Finding all significant de Bruijn subgraphs for bacterial GWAS

**DOI:** 10.1101/2021.11.05.467462

**Authors:** Hector Roux de Bézieux, Leandro Lima, Fanny Perraudeau, Arnaud Mary, Sandrine Dudoit, Laurent Jacob

## Abstract

Genome wide association studies (GWAS), aiming to find genetic variants associated with a trait, have widely been used on bacteria to identify genetic determinants of drug resistance or hypervirulence. Recent bacterial GWAS methods usually rely on *k*-mers, whose presence in a genome can denote variants ranging from single nucleotide polymorphisms to mobile genetic elements. Since many bacterial species include genes that are not shared among all strains, this approach avoids the reliance on a common reference genome. However, the same gene can exist in slightly different versions across different strains, leading to diluted effects when trying to detect its association to a phenotype through *k*-mer based GWAS. Here we propose to overcome this by testing covariates built from closed connected subgraphs of the De Bruijn graph defined over genomic *k*-mers. These covariates are able to capture polymorphic genes as a single entity, improving *k*-mer based GWAS in terms of power and interpretability. As the number of subgraphs is exponential in the number of nodes in the DBG, a method naively testing all possible subgraphs would result in very low statistical power due to multiple testing corrections, and the mere exploration of these subgraphs would quickly become computationally intractable. The concept of testable hypothesis has successfully been used to address both problems in similar contexts. We leverage this concept to test all closed connected subgraphs by proposing a novel enumeration scheme for these objects which fully exploits the pruning opportunity offered by testability, resulting in drastic improvements in computational efficiency. We illustrate this on both real and simulated datasets and also demonstrate how considering subgraphs leads to a more powerful and interpretable method. Our method integrates with existing visual tools to facilitate interpretation. We also provide an implementation of our method, as well as code to reproduce all results at https://github.com/HectorRDB/Caldera_Recomb.

## 1 Introduction

Genome-wide association studies (GWAS) look for genetic variants whose presence or absence is associated with a trait of interest, such as the risk for a person to develop a disease, or the yield for a crop. They were originally used on human genomes using single nucleotide polymorphisms (SNPs) as genetic variants [Visscher et al., 2017]. While SNPs do capture most of the genetic variation in genomes that are *similar enough*, they can miss essential variants in other situations. For example, some bacterial species are known to have large accessory genomes, *i.e*., sets of genes that are not present in every strain in the species. In spite of their name, some of these accessory genes play a central role for some traits of interest, such as antibiotic resistance. In *P. aeruginosa*, for instance, they account for 70% of known genetic determinants of resistance to amikacin [Jaillard et al., 2017]. In this context, *k*-mers—defined as all words of length *k* found in the genomes—have emerged as a popular alternative to SNPs to describe genetic diversity [Sheppard et al., 2013, Earle et al., 2016]. More specifically, bacterial GWAS often test the association between the trait of interest and the presence/absence of *k*-mers. A broad variety of genetic variants–ranging from SNPs to mobile genetic elements or translocations–cause the mutated strains to contain one or several specific *k*-mers. These GWAS are therefore able to capture any of these variants without requiring their prior identification or even definition. On the other hand, *k*-mer-based GWAS suffer from two important limitations. First, interpreting their result is notoriously tedious: any given *k*-mer can belong to several regions of the same genome, and conversely a gene causing the trait of interest can contain a large number of specific *k*-mers. Second, because a resistance-causing gene often exists in slightly different version, its *k*-mers are only present in a fraction of the resistant strains. As a consequence, these *k*-mers are less strongly associated with resistance than the gene itself.

Jaillard et al. [2018] proposed `DBGWAS` to help interpret the result of *k*-mer based GWAS using the De Bruijn graph (DBG [de Bruijn, 1946, Pevzner et al., 2001]), which connects overlapping *k*-mers. Several significant *k*-mers arising from a single polymorphic gene typically aggregate into a somewhat linear subgraph of the DBG (Figure 1), making their interpretation easier. However, `DBGWAS` still tests the individual *k*-mers of this subgraph separately, at the risk of missing causal genes whose presence is too diluted across different versions and therefore different *k*-mers.

**Figure 1:**
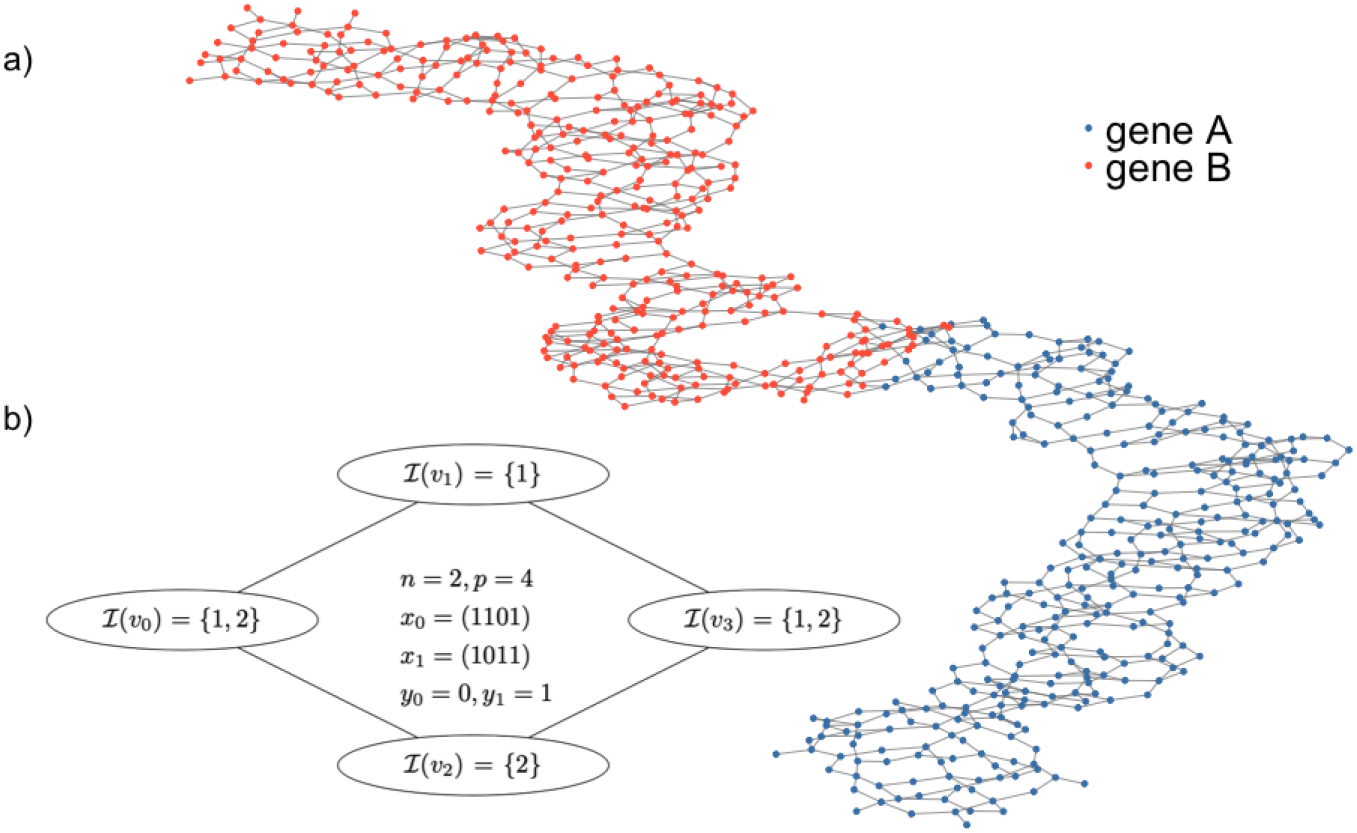
Example of de Bruijn Graphs. **a**. A general example with two genes, each with some variability, resulting in a mostly linear sequence only at the coarse level. More details in the result section. **b**. A simpler setting with 2 samples and 4 nodes. We have three CCSs: {*v*_1_}, {*v*_2_} and {*v*_0_, *v*_1_, *v*_2_, *v*_3_}.

Here we propose to test the association between the phenotype and a single covariate capturing the presence of any version of a gene—or any other potential genetic determinant. Concretely, this covariate indicates the presence of any *k*-mer among those represented in a connected subgraph of the DBG. More specifically, we choose only closed connected subgraphs (CCSs). A CCS is a connected subgraph such that adding any neighbor does not affect the created covariate. Non-closed subgraphs are represented by the same covariate as their closure, and are therefore redundant.

As any such subgraph may represent a causal variant that exists in several version in the dataset, we take an agnostic approach and test the association between the phenotype and one covariate for each connected subgraphs of the DBG. By contrast, `DBGWAS` relies on one covariate for each node of the DBG. This new approach has two potential issues: (1) the number of CCSs grows exponentially with the number of nodes in the DBG, making the task computationally intractable, and (2) adjusting for multiple testing over this very large number of tests leaves little to no power to detect associations. Our method addresses these two issues by using the concept of testability introduced by Tarone [1990]. Tarone’s procedure controls the family-wise error rate (FWER) while disregarding a large number of *non-testable* hypotheses in its multiple testing correction. Intuitively, a covariate representing the presence of any *k*-mer among a growing set that corresponds to larger and larger CCS quickly becomes true for all samples. It thus cannot possibly be associated to any phenotype and can therefore be discarded without being tested or counted towards multiple testing correction. Testability provides a well-grounded and quantitative version of this intuition. Furthermore, since adding nodes to a connected subgraph can only increase the number of present *k*-mers in the corresponding covariate, we can develop a method that rapidly prunes non-testable CCSs, thereby solving the computational problem.

Testability has been used in similar situations, but most existing procedures are restricted to complete [Terada et al., 2013, Minato et al., 2014] or linear graphs [Llinares-López et al., 2015, 2017]. Sese et al. [2014] described an algorithm to test all CCSs: their algorithm combined the testability-based procedure `LAMP` of Terada et al. [2013] with `COIN` [Sese et al., 2010], an enumeration method for CCSs. While no experiment was provided in Sese et al. [2014], we found that a version of this algorithm using an improved version of `LAMP` [Minato et al., 2014, Llinares-López et al., 2015] could find all significant CCSs in graphs with up to 20,000 nodes in less than a day in only the most favorable settings. However the DBG built for typical bacterial GWAS involve millions of nodes, so a more scalable method is necessary to make CCSs testing amenable.

### Our contributions are the following

We introduce a novel, provably complete and non-redundant enumeration scheme for CCSs called `CALDERA`. We also improve an existing pruning criterion for the Cochran-Mantel-Haenszel test. We show that combining these contributions with Tarone’s testability-based procedure makes it possible to find all significant CCSs in a large graph, making it suited to bacterial GWAS. We provide the first implementation of a procedure finding all significant CCSs, along with a user-friendly visualization tool derived from `DBGWAS`. Finally, we demonstrate the advantages of `CALDERA` over competing methods on both simulated and real examples in term of computational speed, statistical power and biological interpretation.

### Notation and goal for CALDERA

We consider a set of *n* samples, 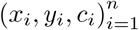, where *x*_*i*_ ∈ {0, 1}^*p*^are *p* binary covariates describing sample *i, y*_*i*_ ∈{0, 1} denotes a binary phenotype, and *c*_*i*_ ∈{1, …, *J*}assigns sample *i* to one population among *J*. We denote *n*_1_ and *n*_2_ the number of samples such that *y*_*i*_ = 0 and 1 respectively. Furthermore, we consider an undirected unweighted connected graph 𝒢 = (𝒱, *E*), where 𝒱 = {*v*_1_, …, *v*_*p*_} and each vertex *v*_*j*_ ∈ 𝒱 is associated with one of the *p* binary covariates represented in *x*. We denote by 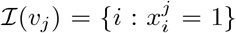. For *i* ∈ [1 : *n*], we note 𝕍_*i*_ = {*v* ∈𝒱 : *i* ∈ℐ (*v*)}. For any connected subgraph 𝒮 = (𝒱′, *E*′), such that 𝒱′⊆𝒱 and *E*′ ⊆ *E*, we let ℐ (𝒮) = ∪*v*∈*S* ℐ (*v*). Of note, this framework addresses both disjunctions and conjunctions, as the latter can simply be obtained by replacing each *x*_*i*_ by its complement. We now properly define the notion of closed connected subgraph and the closure operation (proof in Supplementary S-1.1).

#### Definition 1.

*A connected subgraph* 𝒮 *is closed if and only if there exists no edge* (*v*_1_, *v*_2_) ∈ *E such that v*_1_ ∈𝒮, *v*_2_ ∉ 𝒮, *and* ℐ (𝒮 ∪ {*v*_2_}) = ℐ (𝒮). *We denote by* 𝒞 *the set of all closed connected subgraphs of* 𝒢.

#### Lemma 1.

*For any connected subgraph* 𝒮 *of* 𝒢, *there exists a unique subgraph* 𝒮′∈ 𝒞 *such that* I(𝒮) = ℐ (𝒮′) *and* 𝒮 ⊆𝒮′, *which we note cl*(𝒮).

Assuming that 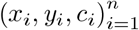 are *n* i.i.d. realizations of random variables **X, Y**, and **C**, our objective is to error rate (FWER, i.e., the chance of at least one Type I error or false positive) at level *α*. Translated in the test null hypotheses of the form 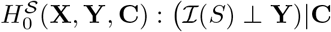 for all 𝒮 ∈ 𝒞, while controlling the family-wise context of GWAS, we want to test the association between the pattern ℐ (𝒮) of each closed connected subgraph 𝒮 with the phenotype **Y**, while controlling for the population structure **C**. We denote 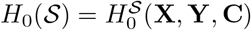 in the remainder of this manuscript, as **X, Y** and **C** are common for all elements of 𝒞.

## 2 Background on significant subgraph detection using testability

Here, we describe the important concept of minimal attainable p-value proposed by Tarone [1990], and how it can be used to (i) retain more power that the Bonferroni procedure while still controlling the FWER and (ii) test more rapidly a large set of hypotheses. Both improvements come from the possibility to discard a large proportion of hypotheses without explicitly testing them and will be exploited in Section 3 to propose our procedure testing all CCSs in 𝒞.

Minimal p-values are a property of discrete tests. For example, Fisher’s exact test [Fisher, 1922] relies on a 2×2 contingency table, whose margins would describe in our case the number of sensitive and resistant bacteria and the number of bacteria whose genome contains or not a genetic variant. Given the margins of this table, only a finite number of cell count values are possible and Fisher’s test can only lead to a finite number of values, the smallest of which is strictly positive (Fig S1). Importantly, this minimal attainable p-value *p*⋆ is entirely determined by the margins of the contingency table: given these margins, *p*⋆ is the minimum over a finite number of possible partitions, and is independent from the actual observed cell counts. Intuitively, strongly imbalanced margins (*e.g*., variants that are present in a very large proportion of samples) cannot possibly lead to small p-values, no matter how the table is filled (*i.e*., how the few samples that do not have the variant are distributed among resistant and sensitive phenotypes).

### 2.1 Using minimal attainable p-values for a tighter FWER control

The family-wise error rate is the probability to incorrectly reject at least one null hypothesis. When testing *N* of them and rejecting those whose p-value *p*_*i*_ is smaller than a threshold 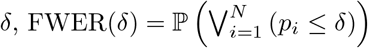, where P is taken over the *N* null distributions 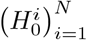. The Bonferroni correction [Bonferroni, 1936] is a common procedure to control the FWER at a level *α*. It is motivated by a simple union bound: as FWER(*δ*) is upper-bounded by 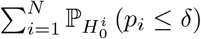 and since by definition 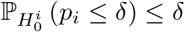, controlling each individual tests at level 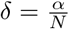 makes the FWER upper-bounded by *α*. Tarone [1990] sharpens this bound, by using the fact that 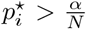 for some hypotheses. Since by definition 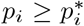, the corresponding term 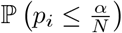 is exactly 0. Therefore, the FWER is actually controlled at level 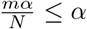 where *m* is the number of testable hypotheses, for which 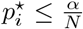. This suggests that using a larger threshold *δ* than the Bonferroni 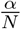 could still control the FWER at level *α*—while rejecting more hypotheses and therefore increasing power. Choosing the largest such *δ* is not a trivial task, as increasing the threshold also decreases the number of non-testable hypotheses. Let *m*(*k*) be the number of testable hypotheses at level 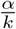, *i.e*. such that 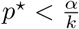. In the worst case, *m*(*k*) = *N* and we recover the Bonferroni procedure. By contrast, we define *k*_0_ as the smallest *k* such that *m*(*k*) ≤ *k*. The largest threshold guaranteeing FWER(*δ*) ≤ α is 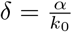.

### 2.2 Using minimal attainable p-values to efficiently explore 𝒞

Provided that enough CCSs have sufficiently large *p*⋆, Tarone’s procedure could therefore address the loss of power incurred when exploring 𝒞. However, naively finding *k*_0_ requires to compute the minimal p-values for all |𝒞|CCSs and iterate through these minimal p-values to adjust the threshold, leaving the computational problem unsolved. A more efficient strategies has been introduced to compute *k*_0_ [Llinares-López et al., 2015, Minato et al., 2014]: starting from *k* = 1 a set ℛ of *testable* hypotheses, *i.e*., of elements with 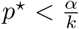 is grown. When |ℛ| becomes larger than *k, k* is incremented to |ℛ|. All hypotheses that are not testable anymore under the new threshold—*i.e*., such that 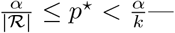 are removed from |ℛ|, and the exploration continues until the point where all testable hypotheses are inℛ and *k* = *k*_0_. This strategy finds *k*_0_ in a single enumeration of all tests, but still requires to compute all minimal p-values, which would not be feasible in our case. However, this search algorithm is also well suited to pruning strategies—a fact already used in [Llinares-López et al., 2015, Minato et al., 2014]. Let *p*⋆(𝒮) be the minimal p-value associated with *H*_0_(𝒮) for a CCS 𝒮. Assuming that for some pairs of subgraphs 𝒮_1_, 𝒮_2_, 𝒮_1_ ⊆𝒮_2_ ⇒ *p*⋆(𝒮_1_) ≤ *p*⋆(𝒮_2_), we can stop exploring all subgraphs including 𝒮_1_ as soon as 𝒮_1_ itself is found non-testable. This monotonicity property is verified when using Fisher’s exact test to test *H*_0_(𝒮): provided that | ℐ (𝒮)| ≥ max(*n*_1_, *n*_2_), *p*⋆ is strictly increasing in | ℐ |, and adding nodes to 𝒮 can only increase | ℐ | (Figure S2). Our main contribution, presented in Section 3 will be an efficient exploration algorithm for 𝒮, which is well suited to pruning.

### 2.3 Controlling for a categorical covariate: the CMH test

When testing for associations, controlling for confounders is essential to avoid spurious discoveries. This is particularly important in bacterial GWAS, where strong population structures can lead to large sets of clade-specific variants to be found associated with a phenotype. The Cochran-Mantel-Haenszel (CMH) test can be used to test associations of two binary variables while controlling for a third categorical variable. It relies on *J* two-by-two association tables such as the one in Table 1, with *j* ∈ {1, …, *J*}, *a*_S,*j*_ = |{*i* : *y*_*i*_ = 1, *i* ∈ℐ (𝒮), *c*_*i*_ = *j*}|, *x*_S,*j*_ = |{*i* : *i* ∈ℐ (𝒮), *c*_*i*_ = *j*}|and *n*_1,*j*_ = |{*i* : *y*_*i*_ = 1, *c*_*i*_ = *j*}.

**Table 1:** Association table in community *j* for subgraph 𝒮, used for the CMH test.

| Variable | $i \in \mathcal{I}(\mathcal{S})$ | $i \notin \mathcal{I}(\mathcal{S})$ | Rows totals |
| --- | --- | --- | --- |
| $\mathbf{y}_i = \mathbf{1}$ | $a_{\mathcal{S},j}$ | $n_{1,j} - a_{\mathcal{S},j}$ | $n_{1,j}$ |
| $\mathbf{y}_i = \mathbf{0}$ | $x_{\mathcal{S},j} - a_{\mathcal{S},j}$ | $n_{2,j} - x_{\mathcal{S},j} + a_{\mathcal{S},j}$ | $n_{2,j}$ |
| Cols Totals | $x_{\mathcal{S},j}$ | $n_j - x_{\mathcal{S},j}$ | $n_j$ |

Like Fisher’s exact test, the CMH test is done conditional on all margins 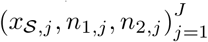. Papaxanthos et al. [2016] furthermore demonstrated that its minimal p-value could be computed in *O*(*J*) (Supplementary S- 1.5) using the margins. However, the minimal p-value of the CMH test does not verify the monotonicity property 𝒮_1_ ⊆𝒮_2_ ⇒ *p*⋆(𝒮_1_) ≤ *p*⋆(𝒮_2_) which is required to prune while exploring 𝒞. Papaxanthos et al. [2016] introduced the envelope, a lower bound on *p*⋆(𝒮), which verifies the monotonicity property. It can also be computed in *O*(*J* log(*J*)) for all S𝒮 such that, for all categories *j, x*_S,*j*_ ≥ max(*n*_1,*j*_, *n*_2,*j*_). This allows for a valid pruning strategy. The condition on *x*_S,*j*_ is the CMH analogous of the;| ℐ (𝒮)| ≥ max(*n*_1_, *n*_2_) condition of Fisher’s test, and can decrease the number of prunable subgraphs as it must be verified for all *J* groups.

## 3 Speeding up the detection of all significant CCSs with CALDERA

We are now ready to present our contributions for scalable detection of significants elements in 𝒞 : an efficient exploration algorithm and an improved envelope for the CMH test, allowing for more pruning in the presence of imbalanced populations.

### 3.1 Critical properties for a fast, Tarone-aware enumeration of 𝒞

We exploit several factors to provide a fast exploration of 𝒞. First, we ensure that it is non-redundant, *i.e*., that each element of is enumerated exactly once, by defining a tree whose nodes are the elements of 𝒞 and propose an algorithm to traverse this tree. Second, the tree is directly built over 𝒞, as opposed to the set of connected subgraphs. The latter option, as proposed in [Sese et al., 2010] is more straightforward to define and to explore and still induces a tree over 𝒞, but yields a much larger object and results in a more expensive traversal. Third, we avoid maintaining subgraph connectivity such as a block-cut tree [Westbrook and Tarjan, 1992]. Such a mechanism is efficient to build a tree over connected subgraphs but is costly to compute. Finally, in order to exploit the pruning opportunity offered by the testing procedure, the exploration should be such that all 𝒮′explored from a given 𝒮 verify 𝒮 ′ ⊃𝒮

Haraguchi et al. [2019], Okuno et al. [2017] define a tree on 𝒞, but the root of the tree corresponds to the entire graph 𝒢: the inclusion relationship along edges of the tree is the opposite to the one we need, making their exploration unsuited to our problem. The `COIN`/`COPINE` algorithm described in Seki and Sese [2008], Sese et al. [2010] builds a tree over the set of connected subgraphs, which induces a tree over 𝒞 but has two drawbacks. First, it maintains an itemtable to enforce a tree structure by avoiding the enumeration of the same element twice. This itemtable has an important memory footprint, and only guarantees a tree structure when exploring in depth first. Secondly, the enumeration of connected subgraphs requires maintaining a list of articulation points along each explored branch, a costly operation.

#### Algorithm 1 Children of 𝒮_*p*_

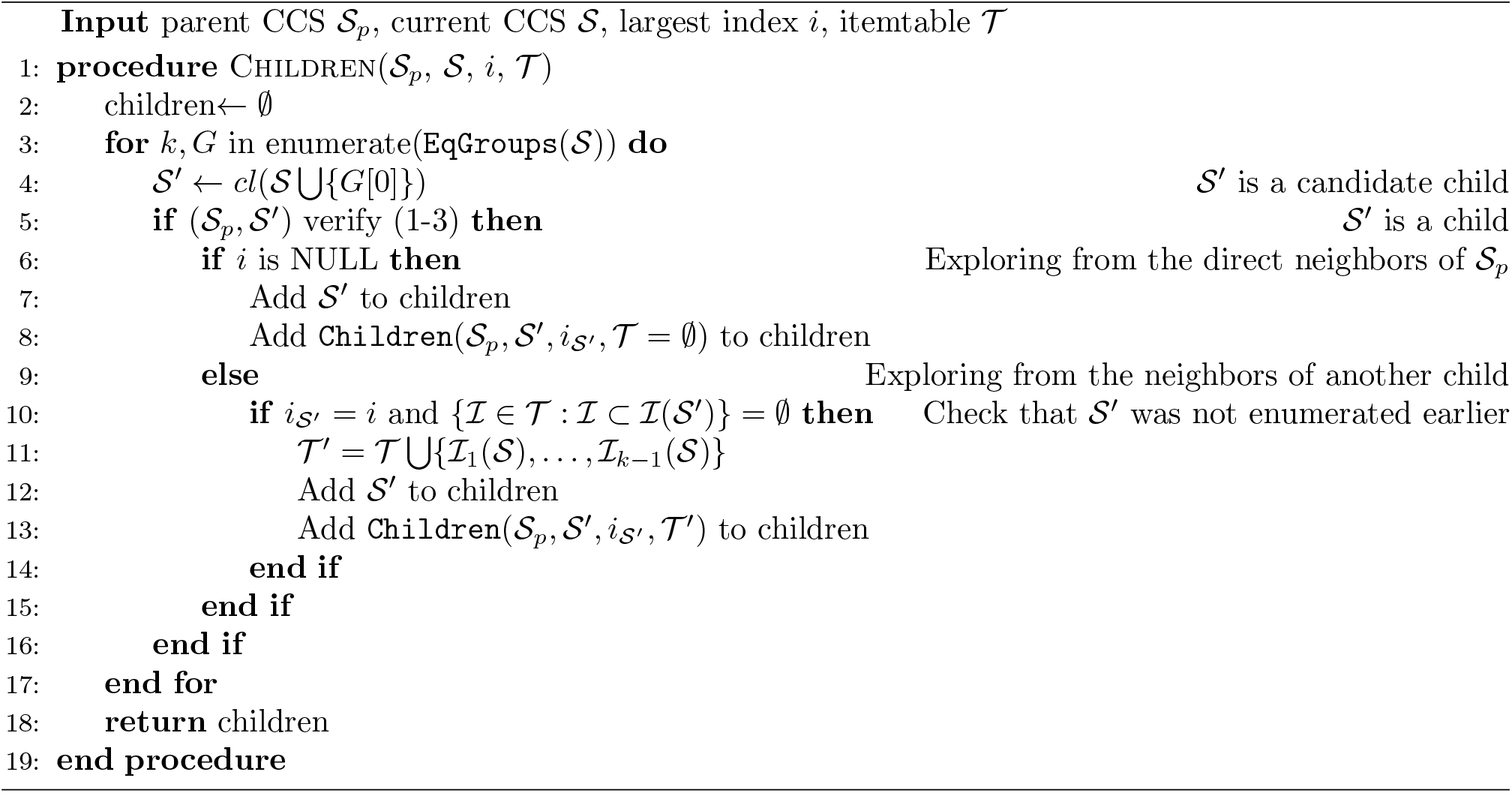

### 3.2 Defining and exploring the tree over 𝒞

In order to build a tree over 𝒞 rooted on the empty CCS, we use a reverse search, introduced in Avis and Fukuda [1993]. Reverse search relies on a reduction operation, which takes one element of the set to be enumerated, and returns a unique, strictly smaller element of the same set. This operation necessarily defines a tree over the elements of the set, by ensuring a unique path between any element and the empty one—the root of the tree. This reduction operation defines the unique parent of every element in the tree. In order to traverse the tree from the root, one needs to inverse the reduction operation, *i.e*. in our setting, given a CCS 𝒮to recover all CCSs that lead to 𝒮by reduction. Here we introduce a reduction operation over 𝒞, as well as n its inversion. We consider the parent operation 𝒫 given by Definition 2 for any element of 𝒞, and show that it defines a valid reduction as introduced above. All proofs are presented in the appendix.

#### Definition 2.

*For a subgraph* 𝒮 ∈𝒞, *we denote* 𝒥 (𝒮) =∩ *v*∈Sℐ(*v*).

- *If* ℐ(𝒮) = 𝒥 (𝒮), *then the parent of* 𝒮 *is* ∅, *i.e*., 𝒫 (𝒮) = ∅.
- *Else we note i*_S_ = max_*i*_{*i* : *i* ∈ℐ (𝒮) \ 𝒥 (𝒮)}. *The parent* 𝒫 (𝒮) *of* 𝒮 *is the connected subgraph of* 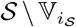 *that contains* 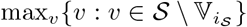.

Note that we arbitrarily assign a number to each node to be able to define the max.

#### Lemma 2.

*The function* 𝒫 *defines a valid reduction over* 𝒞.

Note that we have 𝒮 ⊃𝒫 (𝒮) for all S so this structure allows pruning. Lemma 3 then provides necessary and sufficient conditions for 𝒮 ′∈ C to be a child of 𝒮 ∈ C. The third condition involves the set of neighbouring nodes of 𝒮, *Ne*(𝒮) = {*v* ∈𝒢 \ 𝒮 : ∃*v*_1_ ∈𝒮, (*v, v*_1_) ∈ *E*}.

#### Lemma 3.

*For*, 𝒮 ′ ∈𝒞 *such that* 𝒮 ⊂𝒮 ′≠ ∅, *we have:* 𝒮 = 𝒫 (𝒮 ′) *if and only if the three following conditions are verified:*

*(C1) i*_S*I*_∉ ℐ (𝒮)

*(C2)* 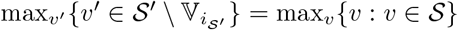

*(C3)* 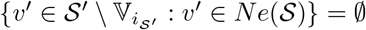, *or written differently*, 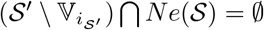.

Using (C1–3) in Lemma 3 to check whether 𝒮 = 𝒫 (𝒮 ′) for any 𝒮^′^does not require to identify the connected components of 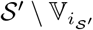, even though the reduction𝒫 itself does rely on these connected components. This property of the inverse reduction is critical for the scalability of `CALDERA`: repeatedly identifying or maintaining these components would be very costly. It is because the reduction operation 𝒫 does not maintain full connectivity: it only retains one of the connected components obtained by removing a subset of its nodes. Doing so comes at a price. Finding all children of 𝒮 is not straightforward, as we must identify and reconnect all the connected components involved—Lemma 3 only provides a way to check if a candidate 𝒮 ′is a child of 𝒮.

More precisely, reducing any CCS 𝒮 ′to its parent 𝒮 involves the removal of a subset 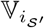 of its nodes, breaking 𝒮 ′ into several connected components—the one containing the largest vertex being retained as the unique parent. For this reason, the reverse search formalized in Algorithm 1 cannot just search for children of 𝒮 among all closures obtained after adding one of its neighbors *Ne*(𝒮) (Lines 6-7): larger CCSs may also lead to 𝒮 by reduction if they involve other connected components that are not in its direct neighborhood. Once a child 𝒮 ′has been identified, we must therefore recursively search for other candidates among the closures obtained after adding one of its neighbors *Ne*(𝒮) (Line 8). This procedure is necessary to reconnect all children that include 𝒮 ′but would leave it as a separated connected component after removing nodes 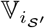. However, Lemma 3 used in Line 5 guarantees only that actual children of 𝒮 are retained, it does not guarantee uniqueness. A redundant exploration would lose the benefit of building a tree over 𝒞 to explore it efficiently. We therefore need an itemtable 𝒯 that keeps track of visited patterns ℐ: if a candidate child 𝒮 ^″^has a pattern ℐ (𝒮 ^″^) that includes the pattern of an already enumerated child from the neighborhood of the same 𝒮 ′, we know that 𝒮 ^″^—and any child that could be obtained from it—has already been visited and the algorithm stops exploring from 𝒮 ^″^. In practice, we do not need to store the full table 𝒯 in order to verify the second condition of Algorithm 1, Line 12. We rely on a concept from [Uno et al., 2004] and further described in Supplementary S-2 to reduce memory footprint.

By Theorem 1, Algorithm 1 solves the problem of inverting the reduction, and therefore of building a tree structure on 𝒞. Of note, Algorithm 1 effectively explores equivalence groups of neighbours yielding the same pattern. Formally, an equivalence group *G*_*k*_(𝒮) ⊂ *Ne*(𝒮) verifies: *v*_1_, *v*_2_ ∈ *G*_*k*_(𝒮) =⇒ ℐ (𝒮 ∪ {*v*_1_}) = ℐ (𝒮 ∪ {*v*_2_}). We name ℐ_*k*_(𝒮) the pattern of the equivalence group *G*_*k*_(𝒮).

#### Theorem 1.

*For any* 𝒮 ∈𝒞, *Algorithm 1 applied on* (𝒮, 𝒮,*NULL*, ∅) *returns the set* {𝒮 ′∈𝒞 : 𝒮 = 𝒫 (𝒮 ′)}.

### 3.3 A breadth-first-search enumeration

We argue that exploring any tree structure on 𝒞 in breadth first will often allow for more pruning than in depth first. At any level, even if the CCSs visited along a branch do increase *k* and therefore lower the testability threshold, all the other CCSs of the level will need to be visited regardless of their testability. By contrast, the increase of *k* gained by visiting all CCSs of the same level in the tree will lower the threshold *α/k* for all CCSs at the next level, making more branches prunable. We demonstrate this in section 4 and provide more intuitive examples in the appendix, (Supplementary S-5.1 and S-5.2). A search in breadth is also easily parallelized since the computation of the minimal p-value, the envelope and the children of every CCS of a given level can be done in parallel, before increasing *k* and updating. ℛBy contrast, a parallelized search in depth-first must share and regularly update *k ℛ* and, which negates the advantages of parallelization.

Algorithm 2 explores 𝒞 through a BFS traversal of the tree defined by the reduction 𝒫, exploiting Algorithm 1 (L.15) to invert the reduction and using this exploration to apply the Tarone testing procedure described in Section 2.2 (L7-12, 14), before finally testing the testable CCSs (L21-25). However, BFS is more memory intensive than DFS (see results). To be able to have a better trade-off between speed and memory, we also implemented a hybrid exploration scheme that is used in practice in which each stage of the tree is explored by batch in a BFS manner, and a DFS is performed over each batch.

### 3.4 Pruning more CCSs when controlling for an imbalanced categorical covariate

The envelope 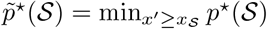 introduced in Papaxanthos et al. [2016] verifies the monotonicity for any subgraph 𝒮 because 𝒮^Ȳ^ ⊇𝒮 ⇒ *x* 𝒮 _*′*_≥ *x* 𝒮. However, the *O*(*J* log *J*) algorithm to compute this envelope only applies to the so called *potentially prunable* subgraphs which are such that *x* 𝒮_,*j*_ ≥ max(*n*_1,*j*_, *n*_2,*j*_) for all subgroups *j* = 1, …, *J* defined by the categorical covariate adjusted for by the CMH test. Pruning can therefore not be done from subgraphs for which at least one of the *J* groups has few occurrences of the corresponding covariate. This limitation arises in Lemma 2 of Papaxanthos et al. [2016], which characterizes the argmin of the envelope of a subgraph S. Lemma 4 lifts this restriction:

#### Lemma 4.

*For any connected subgraph* 𝒮, *the envelope* 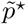 *is attained for an optimum* 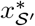 *such that*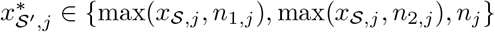.

#### Algorithm 2 List significant closed connected subgraphs

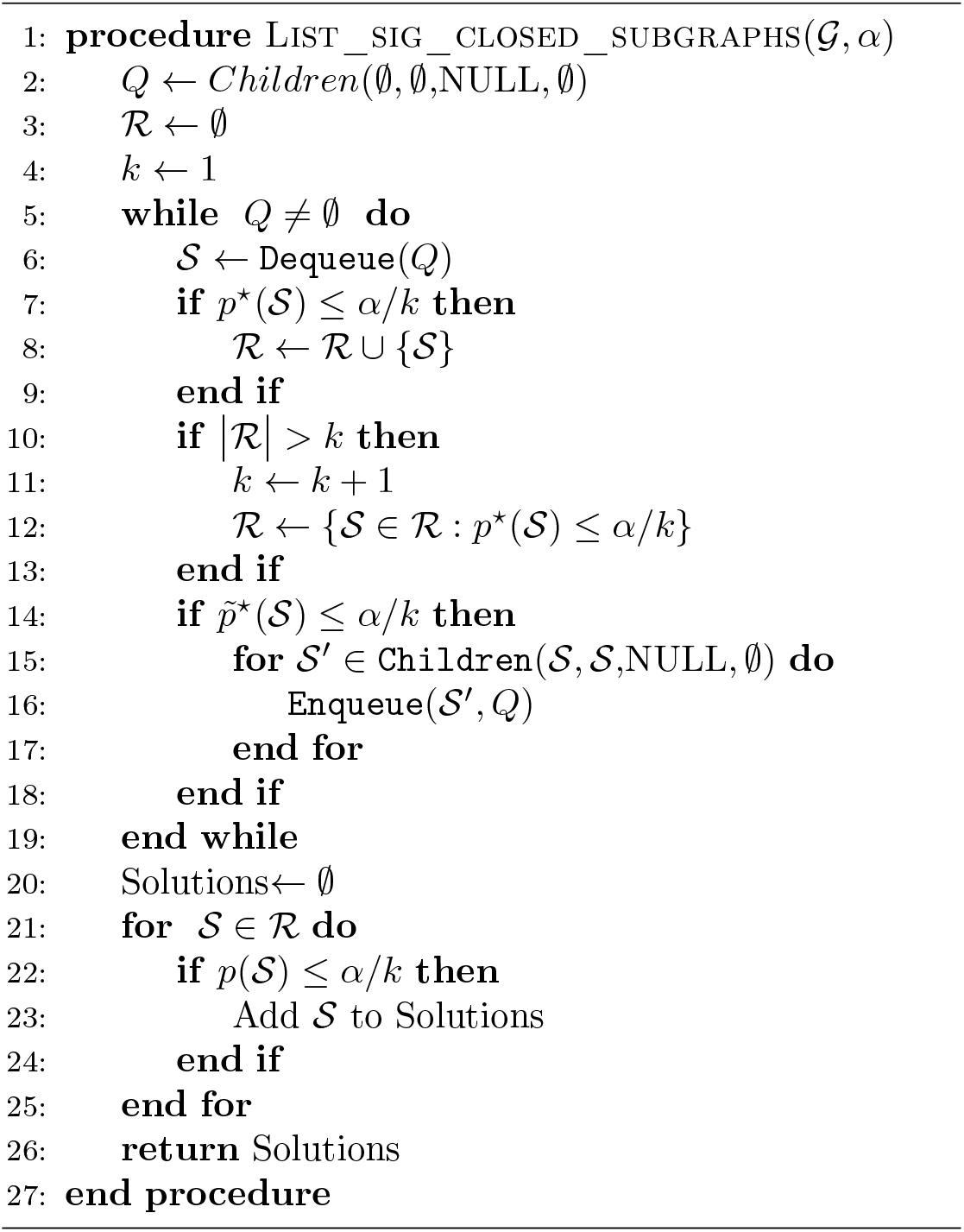

The proof is provided in Supplementary Material S-1.5. Lemma 4 exploits a cruder bound for groups that are not in the increasing regime of the minimal p-value. It recovers the Lemma 2 of Papaxanthos et al. [2016] for potentially prunable subgraphs, while offering an additional pruning opportunity for the other ones. If a subgraph was not potentially prunable only because it was missing the *x*𝒮_,*j*_ ≥max(*n*_1,*j*_, *n*_2,*j*_) condition for one small group *j*, it may still be actually prunable since small groups of samples only affect the CMH test statistic marginally. On the other hand if the condition is not verified for a large group or several small ones, the resulting envelope will be very lose and will not allow for pruning in practice. We provide some intuition in the appendix (Figure S3).

## 4 Experiments

We demonstrate the superiority of `CALDERA` in terms of computational speed, statistical power and biological interpretation. To do so, we rely on both simulated and real datasets.

### 4.1 Datasets and settings

To test the speed of the methods, we generate datasets with *n* samples represented by *p* ∈[100 : 20, 000] covariates, and a graph connecting these covariates. We vary both the proportion *prop* of samples that are resistant, i.e. have a phenotype of 1, and the number of samples. We also perform exploration when changing the value of *α*, which impacts pruning. This generates 4 scenarios to compare the runtimes of the methods, named **Speed 1** to **Speed 4**. More details on implementations and parameters can be found in section S-6. We can also add a binary confounding variable, with *n* = 100 and *p* = 3, 000, where we vary the ratio between the size of those two populations, *n*_2_*/n*_1_. This is useful to test the speed gains provided by the new lower bound provided in `CALDERA`. This scenario is called **Imbalance**

To test the power of the different methods, we rely on a simulation where the ground truth is known, named **Exploration**. We generate a dataset with *n* = 100 samples, 50 of each phenotype, where two genes A and B are present. Gene A is present for all samples while gene B is only present for resistant samples. We introduce heterogeneity such that the DBG of the two genes is only linear at a coarse level (Fig.1b). More details for the setting of those simulations are provided in S-7.

We also rely on two real datasets. The first, which we name **Pseudomonas**, consists of the *n* = 280 *Pseudomonas Aeruginosa* genomes along with their resistance phenotype to amikacin, used in `DBGWAS` [Jaillard et al., 2018]. The bacteria are partitioned based on k-mean into two distinct groups. The compacted DBG is constructed using the *k*-mers with *k* = 31 (default) using `DBGWAS`, leading to a graph with over 2.3 million nodes and average degree ∼2.7. The second, named **Akkermansia**, consists of the Akkermansia muciniphila genomes collected in Karcher et al. [2021]. We use host information as covariates: we want to identify genetic sequences that are associated with a body mass index (BMI) over 30. The DBG constructed over those *n* = 401 strains has 1.3 million nodes with an average degree of∼ 2.7. On these two real datasets, we rely on heuristics to choose the level *α* at which the FWER is controlled and the number of stages explored in the BFS search—a full exploration being too memory intensive. The level is fixed at the lowest value at which 10 CCSs at stage 1 of the BFS (*i.e*, unitig closures) are found significant. The stage is chosen by stopping when the number of unitigs covered by a significant CCS reaches a plateau—suggesting that further exploration would not bring much novelty.

### 4.2 Speed gains of CALDERA

`COIN` [Sese et al., 2014] was to our knowledge the only described algorithm to identify significant CCSs, combining the enumeration method of `COPINE` with the `LAMP` algorithm. Minato et al. [2014] presented a provably superior version of `LAMP`, which we denote `LAMP2`. Since no implementation was provided in Sese et al. [2014], we implemented as a baseline `COIN+LAMP2`. Since `CALDERA` and `COIN+LAMP2` both rely on the same statistical procedures (the identification of testable hypotheses with Fisher’s test), the set of significant CCSs found is the same regardless of the method. In addition to `COIN+LAMP2`, we benchmark 3 versions of `CALDERA`. The first one, closest to `COIN+LAMP2`, is the DFS implementation. The second one is the BFS implementation, where we modify the enumeration order of the elements of 𝒞 to promote pruning. The last is a parallelized BFS implementation, using 5 cores.

#### Benefit of CALDERA ’s exploration scheme

In Figure 2a, representing the results of **Speed 2**, we see that the ranking in speed is uniform over all value of *p*, with `COIN+LAMP2` being the slowest, followed by the DFS and BFS implementation, and finally the parallelized version of `CALDERA`. For *p* = 20, 000, `COIN+LAMP2` runs in 2h20 while the parallelized version of `CALDERA` takes 5 minutes. The ranking is the same for **Speed 1, Speed 3** and **Speed 4** (see Supplementary S-6). For example, for **Speed 1** and *p* = 20, 000, `COIN+LAMP2` times out (two day threshold) before finishing while the parallelized version of `CALDERA` runs in 6 hours. Over all parameter values, the average ratio of runtime for `COIN+LAMP2` over `CALDERA BFS` with 5 cores is 76 and we tested `CALDERA` on values of *p* = 100, 000 in 14h. More details on memory usage can be found in section S-6.

**Figure 2:**
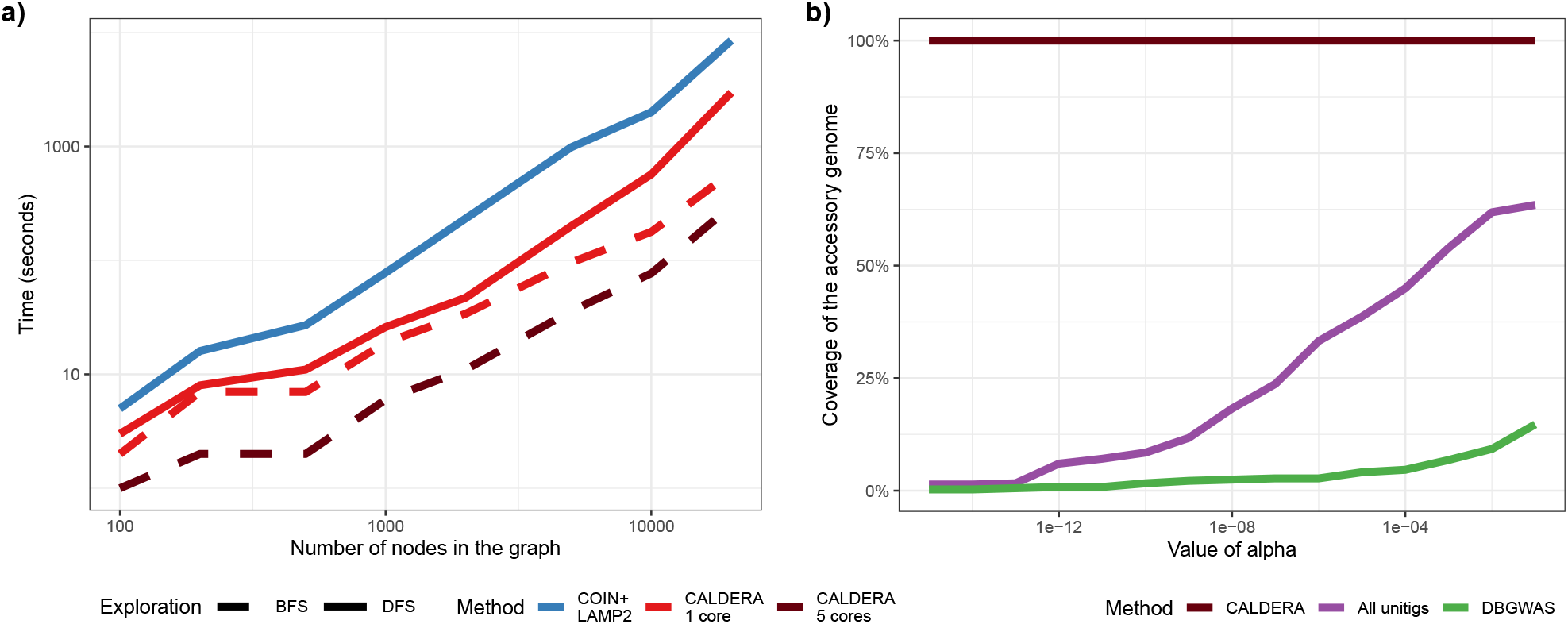
Results of CALDERA. **a**. Run times for `CALDERA` and `COIN+LAMP` on graphs with various values of *p*. In this setting, *n* = 100. **b**. Proportion of all unitigs associated with the resistant phenotype that are found to be significant by `CALDERA`, the `LAMP2` procedure on all unitigs and `DBGWAS`, as the value of *α* changes.

On the larger **Pseudomonas** dataset, even `CALDERA` is unable to explore the entire 𝒞 with the heuristic level *α* = 10^−6^, but we observe that the unitigs covered by the significant CCSs reached a plateau after the first 6 stages of the BFS. `CALDERA` took 3h20 on 4 cores, using 200Gb of RAM to complete these stages using batches of size 200, 000, leading to *k*_0_ = 4, 671, 265 potentially testable CCSs and only 39 significant ones. For comparison, after running for 24h, `COIN+LAMP2` was exploring the tree structure with a running *k*_0_ = 10^5^. We provide a more general analysis of the computational cost of `CALDERA` against the number of BFS stages in Supplementary S-9 and recommend using a similar analysis and stop after a few stages in cases where a full exploration is no feasible.

#### Benefit of CALDERA’s lower-bound on runtime for imbalanced population

For extreme ratios— below 0.02—the new lower bound allows much more pruning and enumerates an order of magnitude fewer elements of 𝒞. Up to a ratio of 0.1, the new lower bound leads to a decrease of at least 10% in the number of explored subgraphs (see Fig S7).

### 4.3 Power gains of CALDERA

As mentioned above, `COIN+LAMP2` and all versions of `CALDERA` rely on the same statistical procedures and therefore find an identical list of significant CCSs for a given level of *α*. However, we can compare the power of `CALDERA` with two other alternatives. `DBGWAS` tests individual unitigs for association with a phenotype, using a mixed-model. We also use the `LAMP2` procedure when testing `All Unitigs` separately using Fisher’s test—like `CALDERA`.

We run all three methods on the dataset **Exploration** and measure how many of the 367 unitigs of gene B are called significant, when controlling the FWER at a varying level *α*. For `CALDERA`, a significant unitig is one that is contained in a significant CCS. Even when controlling the FWER at very low levels (*α* = 10^−16^), `CALDERA` correctly recovers the entirety of the resistant gene. On the other hand, the other two methods fail to ever recover the entire gene, even at *α* = 0.1. This clearly show the enhanced power of `CALDERA`: because of variations along the genome, the association of any individual unitig with the phenotype is weak, while a covariate that jointly represents all 367 unitigs of the resistant gene is very strongly associated with that phenotype.

We also apply those three methods to the **Pseudomonas** dataset. While there is no ground truth, this dataset contains two confirmed genetic variants linked to resistance to amikacin: a SNP on the aac(6’) gene, represented by one unitig, and the pHS87b plasmid, represented by 476 unitigs. This allows us to see how the methods handle those different scales. At the default *α* = 10^−6^, `CALDERA` find the aac(6’) mutation as one CCS, and finds significant CCSs that covers 96% of the plasmid. Those two components represent 59% of all significant unitigs. In contrast, `All Unitigs` and `DBGWAS` do recover the mutation but only 34% and 0% of the plasmid respectively. Even at *α* = 0.1, `All Unitigs` and `DBGWAS` only recover respectively 72% and 8% of the plasmid. Moreover, while it is not possible to compute a false negative rate on real dataset, we can see that, at this level, the two known sequences—the plasmid and the aac(6’) mutation—only represent 6% and 17% of all significant unitigs.

### 4.4 Simplified biological interpretation

Biological interpretation in `DBGWAS` or `CALDERA` happens at the component levels: significant unitigs or CCSs separated by only a a few non-significant unitigs are displayed as one component. Unitigs can also be annotated using various databases, to enhance interpretation. Components are ranked in order of decreasing p-values, choosing the smallest p-value among all unitigs/CCSs. As such, both the number of components and their rankings will impact the ease of interpretation.

Figure 3 gives two examples. On panel **a)**, we see the results of running `CALDERA` on the **Akkermansia** dataset. Only one CCS is significant, while `DBGWAS` returns no significant unitig. All the unitigs in the significant CCS (colored in green) are annotated, using the *RefSeq* database of all known Akkermansia muciniphila proteins as a reference [Tatusova et al., 2016], and map to a common gene. This gene is not well annotated (hypothetical protein) but partially map to a Tubulin / FtsZ_GTPase Akkermansia muciniphila protein.

**Figure 3:**
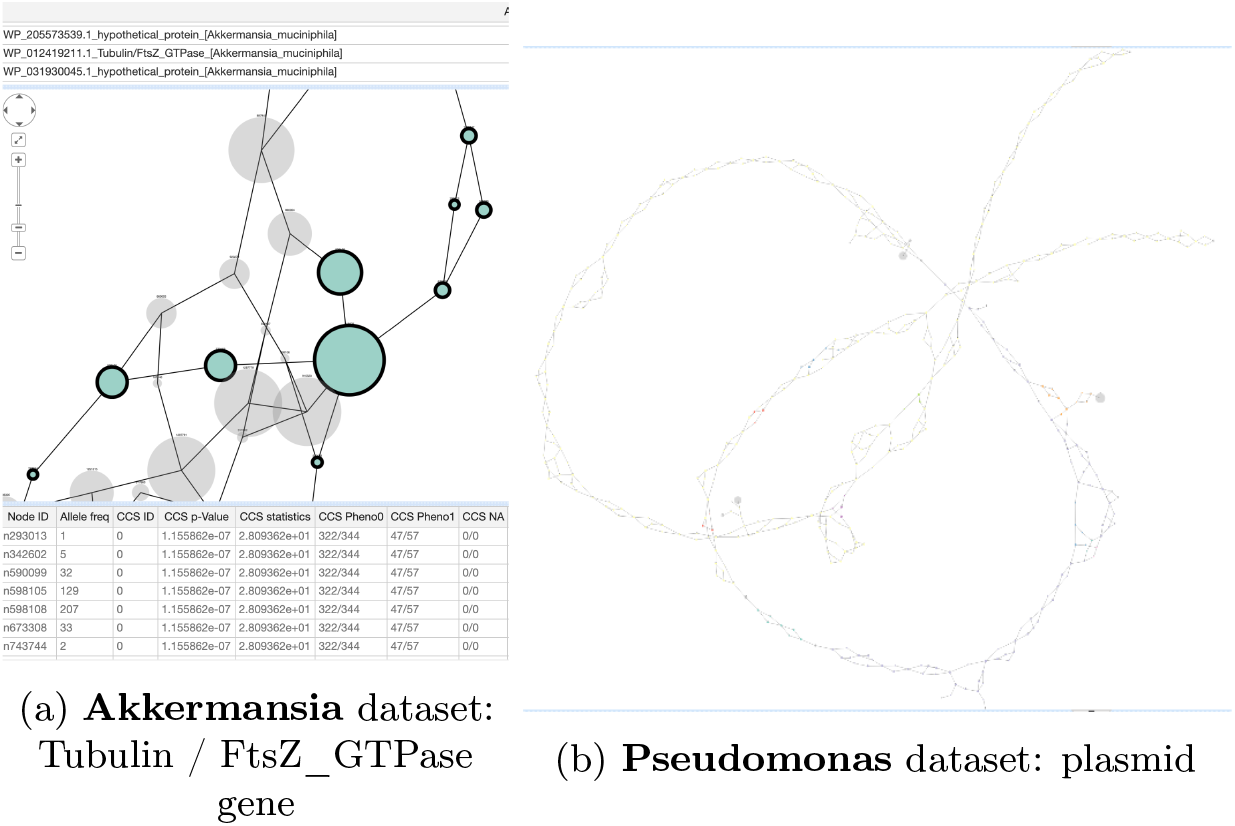
Screenshot from the output of `CALDERA`. We select the first component, that is the one which contains the most significant CCS. Unitigs belonging to the same CCS are colored in the same way. If two significant CCSs are less than two unitigs apart, they are represented in the same component.

On panel **b)**, we see the plasmid, returned as the first component for `CALDERA`. Visually, we can see clearly a broad circular graph, with local genetic variations. On the **Pseudomonas** dataset, `CALDERA` returns 8 components: the first is the entire plasmid, returned as one component. The second is the aac(6’) mutation. `DBGWAS` always ranks the aac(6’) mutation first but never returns the plasmid as one component, even when controlling the FWER at a level of 0.1 (3 components, the first one ranked fourth). Moreover, at this level, `DBWGAS` returns 77 components, making the interpretation much harder.

## 5 Discussion

This article presented `CALDERA`, an algorithm to enumerate all significant closed connected subgraphs. `CALDERA` easily scales to large datasets, relying on an efficient structure on 𝒞 and an exploration scheme that leverages the pruning opportunity offered by discrete statistics. This increased computational speed allows to deploy this method to De Bruijn graph-based bacterial GWAS, which we demonstrate on two real examples. Moreover, we show that considering the CCSs, as done by `CALDERA`, leads to increased power and facilitates interpretation, compared to previous methods to performed statistical tests at the node level. `CALDERA` can better detect low signal caused by variability in genetic elements. It also returns larger and more coherent outputs that are easier to interpret.

We extensively discussed how `CALDERA` performs on bacterial GWAS. However, `CALDERA` can also be used for other tests of association involving a graph structure. We provide in S-8 an example: we look at the association between SNPs on *A. thaliana* genomes and a “date to flowering” phenotype. In that setting, the graph is a regulatory network on the genes and the objective is to identify subnetworks whose disruption by at least one mutation is associated with the phenotype.

In settings where the node is a more natural object than the CCS, discrete testing can still be used to take advantage of Tarone [1990]’s procedure and increase power. However, pruning will no longer be possible, unless some other order can be established between nodes that preserve the order of minimal p-values.

For now, `CALDERA` does not scale to datasets with hundreds of millions of nodes that are possible in metagenome-wide association studies. Future work that focuses on incorporating pre-processing schemes before `CALDERA` would be needed to compact the graph to both reduce its size and facilitate pruning by increasing the average |ℐ (*v*_*j*_)|.

## Acknowledgment

LJ is funded by the Agence Nationale de la Recherche ANR-17-CE23-0011-01 (FAST-BIG). A part of this work was performed using the computing facilities of the CC LBBE/PRABI.

## S-1 Proofs

### S-1.1 Lemma 1: correctness of the closure

Lemma 1 provides that the operator *cl* is well defined on connected subgraphs.

**Proof of Lemma 1** First let’s show that there exists 𝒮′ ∈ 𝒞 such that ℐ(𝒮) = ℐ(𝒮′) and 𝒮 ⊆ 𝒮′. Let 𝒮′ be a (inclusionwise) maximal connected subgraph containing 𝒮 and such that ℐ(𝒮) = ℐ(𝒮′). By maximality of 𝒮′, for every edge (*v*_1_, *v*_2_) ∈ *E* with *v*_1_ ∈ 𝒮′ and *v*_2_ ∉ 𝒮′, we have ℐ(𝒮′ ∪ {*v*_2_}) ≠ ℐ(𝒮) = ℐ(𝒮′), thus 𝒮′ ∈ 𝒞.

Now let’s show that such a subgraph is unique. Assume that there exits two different subgraphs 𝒮_1_ and 𝒮_2_in 𝒞 such that 𝒮 ⊆ 𝒮_1_ and 𝒮 ⊆ *𝒮*_2_ with ℐ(𝒮) = ℐ(𝒮_1_) = ℐ(𝒮_2_). Since 𝒮_1_≠ 𝒮_2_, at least one of the subgraphs 𝒮_1_ \ 𝒮_2_ and 𝒮_2_ \ 𝒮_1_ is not empty. Assume without loss of generality that 𝒮_1_ \ 𝒮_2_≠ ∅. Since 𝒮_1_ is connected and since 𝒮_1_ ∩ 𝒮_2_ ⊇ 𝒮 ≠ ∅, there is at least one edge (*u, v*) with *u* ∈ 𝒮_1_ ∩ 𝒮_2_ and *v* ∈ 𝒮_1_ \ 𝒮_2_. This leads to a contradiction since the edge (*u, v*) is such that *u* ∈ 𝒮_2_, *v* ∉ 𝒮_2_ and ℐ(𝒮_2_ ∪ *v*) = ℐ(𝒮) = ℐ(𝒮_2_), which is in contradiction with 𝒮_2_ ∈ 𝒞.

### S-1.2 Lemma 2: 𝒫 is a valid reduction

**Case if** ℐ(𝒮) = 𝒥 (𝒮): Then, either 𝒮 = ∅ which has trivially no parent by this reduction. Or all nodes of 𝒮 contain exactly the same pattern. For any *v* ∈ 𝒮, 𝒮 = *cl*(*v*). 𝒮 is a root of our exploration. Its parent is ∅ ⊆ 𝒮. Note that, to avoid enumerating those roots more than once, we only start from *v*_max_ = max 𝒮.

**Case if** *i*_𝒮_ **is defined:** Then, *i*_𝒮_ ∈ ℐ(𝒮) so 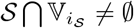 and *i*_𝒮_ ∉ ℐ(𝒮) so 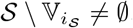. Therefore, there is at least one connected component in 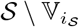. Moreover, any connected component of 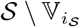 is included but not equal to 𝒮. From [Haraguchi et al., 2019], Lemma 1, we know that, if 𝒮 ∈ 𝒞, any connected component of 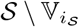 is also in 𝒞. So any connected component of 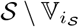 can be defined as a parent of 𝒮. To identify a unique parent, we select 𝒮_*p*_, the one with the highest node number, 𝒮_*p*_. Since 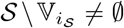 and 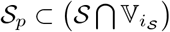, then 𝒮_*p*_ ⊊ 𝒮: it is a strictly smaller subgraph by inclusion. This proves that reduction defines a unique parent and proves that the reduction is valid.

### S-1.3 Lemma 3: conditions (C1-3) are necessary and sufficient for 𝒮 = 𝒫(𝒮*′*)

We first prove the following lemma:

#### Lemma 5.

*For two subgraphs* 𝒮_1_, 𝒮_2_ ∈ 𝒞, *if* 𝒮_1_ ⊂ 𝒮_2_, *then cl*(𝒮_1_) ⊂ *cl*(𝒮_2_).

**Proof:** Since 𝒮_1_ ⊂ *cl*(𝒮_1_) and 𝒮_1_ ⊂ 𝒮_2_ ⊂ *cl*(𝒮_2_), we then known that *cl*(𝒮_1_) ∩ *cl* (𝒮_2_) ≠ ∅. Let’ now assume that *cl*(𝒮_1_) ⊈ *cl*(𝒮_2_). Since *cl*(𝒮_1_) is a connected subgraph, there exists *v* ∈ *cl*(𝒮_1_) ≠ *Ne*(*cl*(𝒮_2_)). *v* ∈ *cl*(𝒮_1_) so ℐ(*v*) ⊂ ℐ(*cl*(𝒮_1_)) = ℐ(𝒮_1_). But 𝒮_1_ ⊂ 𝒮_2_ so ℐ(𝒮_1_)] ⊂ ℐ(𝒮_2_). Therefore, ℐ(*v*) ⊂ ℐ(*cl*(𝒮_2_)) and *v* ∈ *Ne*(*cl*(𝒮_2_)). This contradicts the definition of the closure. So we prove the lemma.

#### S-1.3.1 Proof that for any 𝒮′, (𝒮 = 𝒫(𝒮′), 𝒮′) verify (*C*1 − 3)

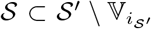 so *i*_𝒮_*′* ∉ ℐ(𝒮). This proves (1). 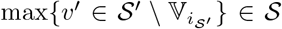 by construction of the parent so max 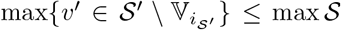. Moreover, 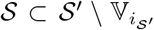 so 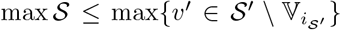. So 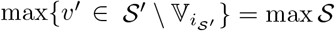, this proves (2).

Suppose (3) is false. Then, there exists 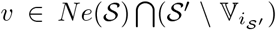 so 𝒮_2_ = *cl*(𝒮⋃{*v*}) ⊂ 𝒮′, 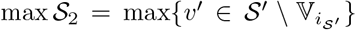 since 𝒮 ⊂ 𝒮_2_ and 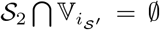 so 𝒮_2_⊂ 𝒫(𝒮′). But 𝒮_2_⊋ 𝒮 = 𝒫(𝒮). This is not possible. So (3) is true.

This proves the implication in the first sense.

#### S-1.3.2 Proof that for any (𝒮, 𝒮′) that verify (*C*1 − 3), 𝒮 = 𝒫(𝒮′)

We consider two closed connected subgraph 𝒮, 𝒮′ ∈ 𝒞 that verify (1-3). We want to prove that 𝒫(𝒮′) = 𝒮. Point (1) insures that 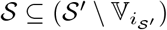. Since 𝒮 ∈ 𝒞 and contains the maximal node (from (2)), this ensures that 𝒮 ⊆ 𝒫(𝒮′).

Suppose 𝒮 𝒫 (𝒮′). Then) 𝒫(𝒮′) \ 𝒮 ≠ ∅. In particular, since 𝒮 and 𝒫(𝒮′) are both connected subgraphs, there exists *v*′ ∈ 𝒫(𝒮′) \ 𝒮 ∩ *Ne*(𝒮). Since this neighbour is in 𝒫(𝒮′), it is also in 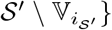}. That is impossible from (3). So 𝒮 = 𝒫 (𝒮′). (Note that point (3) includes the fact that *i*_𝒮_*′* ∈ ℐ (*v*)).

This proves the converse implication.

### S-1.4 Theorem 1: Algorithm 1 correctly inverts the reduction

We consider a subgraph 𝒮′ and its parent 𝒮 = 𝒫(𝒮′).

We first show two lemmas

#### Lemma 6.

*For two subgraphs* 𝒮_1_, 𝒮_2_ ∈ 𝒞, *if* 𝒮_1_ ⊂ 𝒮_2_, *then* 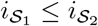.

**Proof:**

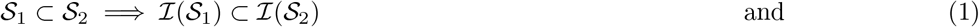

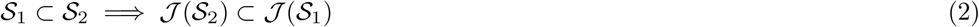

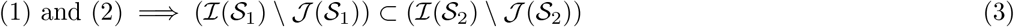

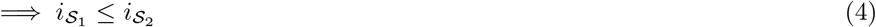

#### Lemma 7.

*For a subgraph* 𝒮′ ∈ 𝒞 *such that* 𝒮 = 𝒫(𝒮′) ≠ ∅, *any subgraph* 𝒮_2_ ∈ 𝒞 *that verifies:*

- 𝒮 ⊊ 𝒮_2_
- 𝒮_2_ ⊂ 𝒮′

*is a child of* 𝒮, *that is* 𝒫(𝒮_2_) = 𝒮

**Proof:** We know that 𝒮 ⊊ 𝒮_2_ so *Ne*(𝒮) ∩ 𝒮_2_ ≠ ∅. Since 𝒮_2_ ⊂ 𝒮′, *Ne*(𝒮) ∩ 𝒮_2_ ⊂ 𝒮′ so, from (3) for 𝒮, 𝒮′, we have 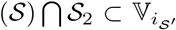. So *i*_𝒮_*′* ∈ ℐ(𝒮_2_). *i*_𝒮_*′* ∉ ℐ(𝒮) so *i*_𝒮_*′* ∉𝒥 (𝒮_2_). Therefore, 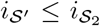. But since 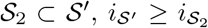. So 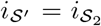. Then, we know that 𝒮, 𝒮_2_ verifies (1). Since 𝒮 ⊊ 𝒮_2_, we also have (2). Finally 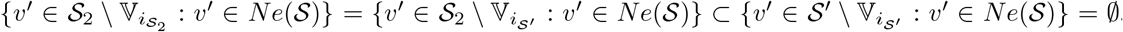. This proves (3). Since we have (1-3), we know that 𝒫(𝒮_2_) = 𝒮.

**Main proof:** Now let us prove the main result: Let’ consider a subgraph 𝒮 and assume that we cannot generate 𝒮′ that verifies 𝒫(𝒮′) = 𝒮 with the procedure from algorithm 1.

We take 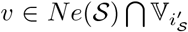. 𝒮_*d*_ = *cl* (𝒮 ⋃ (*v*}) verifies 𝒮_*d*_ ∈ 𝒞, 𝒮 ⊆ 𝒮_*d*_ and 𝒮_*d*_ ⊂ 𝒮′ (from Lemma 5). So, from lemma 7, we know that 𝒫(𝒮_*d*_) = 𝒮 so (𝒮, 𝒮_*d*_) verifies (1-3). Therefore, we generate _*d*_ through lines 4-8 of algorithm 1. (Note that we can call 𝒮_*d*_ a *direct* child of 𝒮, and the other subgraphs generated via lines 11-15 as its siblings).

We know that we generate through algorithm 1 from 𝒮_*d*_ a set of subgraphs 𝒮_*s*_ ⊂ 𝒮′. Let’s then consider the largest 𝒮″ ⊊ 𝒮′ generated with the algorithm 1, that is the one with the largest number of nodes. We note 𝒯 the itemtable that accompanies the creation of 𝒮″. Note that we know that *i*_𝒮_*″* = *i*_𝒮_*′* since *i*_𝒮_*″* ≤ *i*_𝒮_*′*. (lemma 6) and 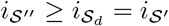.

By assumption, 𝒮″ ⊊ 𝒮′. Therefore, there exists a neighbour *v* ∈ *Ne*(𝒮″) ∩ 𝒮′ since 𝒮′ and 𝒮″ are connected subgraphs. Moreover, 𝒮_2_ = *cl*(𝒮^″^⋃{*v*})) is a child of 𝒮 by Lemma 7 so (𝒮, verify (1-3) by Lemma 2.

**Case 1:** {ℐ ∈ 𝒯 : ℐ ⊂ ℐ(𝒮)2)} = ∅: Since 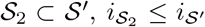. However, since 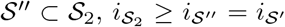. So 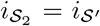. Moreover, we already know that (𝒮, 𝒮_2_) verify (1-3). We therefore try 𝒮_2_ following Line 5, we checked all the conditions (Line 11-12). That means we can created 𝒮_2_ ⊋ 𝒮″ which contradicts our assumption that 𝒮″ is the largest closed subgraph strictly included in 𝒮′ that could be generated.

**Case 2:** {ℐ ∈ 𝒯 : ℐ ⊂ ℐ (𝒮)2)} ≠ ∅ : We note *v*_1_, …, *v*_*l*_ the sequence that created 𝒮″ from 𝒮_*d*_ through successive additions and closures (following Line 5). At one point in that process, we added a pattern to that is now contained in ℐ(*cl*(𝒮^″^ ⋃ {*v*})), let’s say when adding *v*_*k*_. Note that it cannot be the same pattern otherwise 𝒮″ would not be closed. This pattern was linked to another equivalence group (Line 13).. If we consider *v*′ a node from that group, we will then construct a subgraph using the sequence *v*_1_, …, *v*_*k*−1_, *v*′, *v*_*k*_, …, …*v*_*l*_. Note that since at each new addition, the constructed graph is included in 𝒮_2_, it’s also included in 𝒮′. Moreover, each one contains 𝒮 and a node from 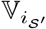 by construction (since it contains 𝒮_*d*_). 𝒮o, using Lemma 2, we know that those additions verifies the conditions of Lines 11 and 12: they are valid additions according to our algorithm. This way, we can create a subgraph that contains 𝒮″ and *v*′ with our procedure. This contradicts our assumption that 𝒮″ is the largest closed subgraph strictly included in 𝒮′ that could be generated.

This proves that all 𝒮′ will be generated from 𝒮 and therefore that we have properly inverted the reduction

### S-1.5 Proof of Lemma 4

The next two lemmas are directly taken from [Papaxanthos et al., 2016].

#### Minimal p-value of the CMH test

##### Lemma 8

([Papaxanthos et al., 2016]). *The minimal p-value of the CMH test can be computed in O*(*J*).

**Proof** The p-value associated with the CHM test, conditioning on the margins of all the tables, is:

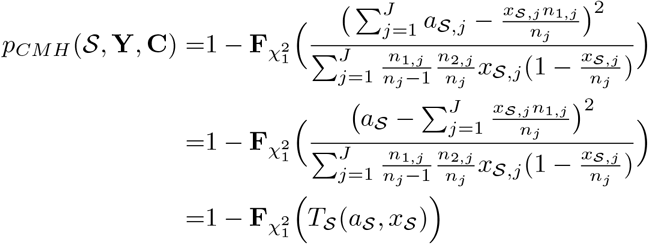

Since 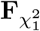 is monotonically increasing, the minimal p-value is obtained for the smallest value *T*_𝒮_ (*a*_𝒮_, *x*_𝒮_. This is a function of *a*_𝒮_ that is quadratic with a positive definite hessian[Papaxanthos et al., 2016] so the function is maximal for min *a*_𝒮_ or max *a*_𝒮_. We have that *a*_𝒮,*j*,min_ = 0 if *x*_𝒮,*j*_ ≤ *n*_2,*j*_ and *a*_𝒮,*j*,min_ = *x*_𝒮,*j*_ − *n*_2,*j*_; and *a*_𝒮,*j*,max_ = *x*_𝒮,*j*_ for *x*_𝒮,*j*_ ≤ *n*_1,*j*_ and *a*_𝒮,*j*,max_ = *n*_1,*j*_ otherwise. So, 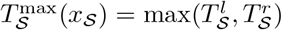 where

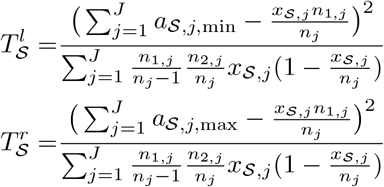

#### Computing the envelope

##### Definition 3.

*For each* 𝒮 ∈ 𝒞, *the envelope of* 𝒮 *is defined as* 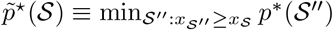.

##### Lemma 9

([Papaxanthos et al., 2016]). *If a subgraph is prunable, i.e* 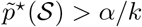, *then any subgraph* 𝒮′ ⊃ 𝒮 *is also prunable*

**Proof** :

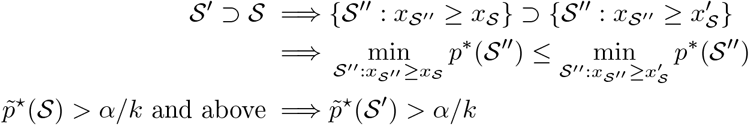

**Proof of Lemma 4** We consider the function

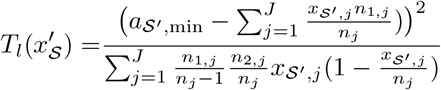

with *x*_𝒮,*j*_ ≤ *x*_𝒮_*′* _,*j*_≤ *n*_*j*_ and we will look at the partial derivatives. We add a few notations:

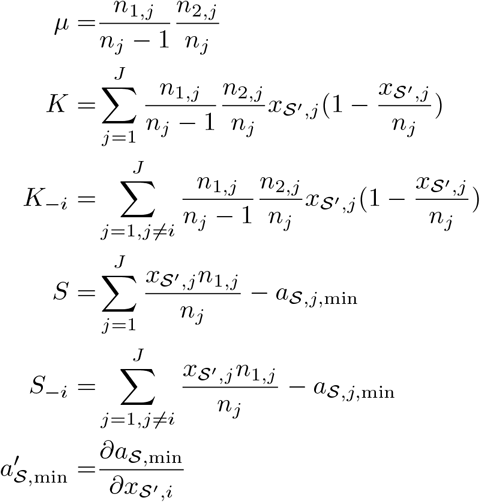

We then have

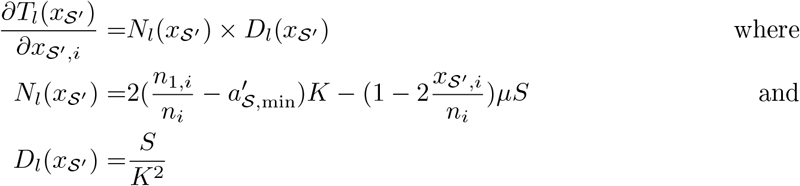

For all 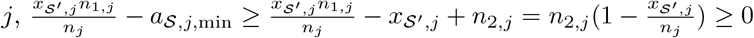 so *Dl* (*x*_𝒮_*′*) ≥ 0. We only need look at *N*_*l*_(*x*_𝒮_*′*) to find maxima. We can also note that consequently, *𝒮* ≥ 0 and *𝒮*_−*i*_ ≥ 0.

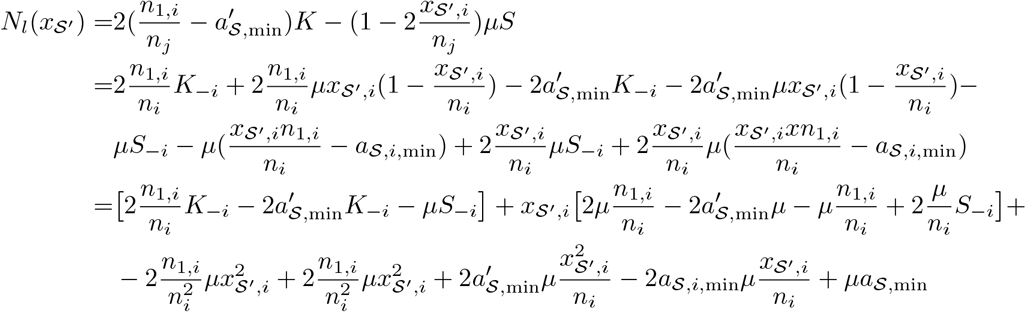

We then have two cases:

*x*_𝒮_*′* _,*i*_≤ *n*_2,*i*_: Then, *a*_𝒮,*i*,min_ = 0 and *a*′ _𝒮,min_ = 0. So

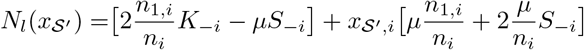

It is an affine function with a positive slope. Moreover, at the boundary at *n*_2,*i*_, the function is positive. So the function is maximal at *n*_2,*i*_.

*x*_𝒮_*′* _,*i*_≥ *n*_2,*i*_: Then, *a*_𝒮,*i*,min_= *x*_𝒮_*′* _,*i*_− *n*_2,*i*_and *a*′_𝒮,min_= 1. So

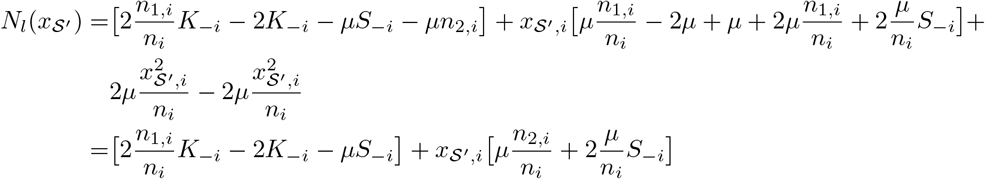

So *N*_*l*_(*x*_𝒮_*′*) is an affine by-piece function of *x*_𝒮_*′* _,*i*_, whose slope *A*_*l*_(*x*_𝒮_*′* _,−*i*_) ≥ 0. So, the only possible maxima are at the boundary, where *x*_𝒮_*′* _,*i*_= *n*_2,*i*_ or *x*_𝒮_*′* _,*i*_= *n*_*i*_. Since this is true for all values of *x*_𝒮_*′* _,−*i*_, we know that we can only achieve a maximum for *T*_*l*_ at the boundaries. So the only two possible maxima are *n*_2,*i*_ and *n*_*i*_. Note that, in the case where *x*_𝒮_*′* _,*i*_≥ *n*_2,*i*_, then the possible maxima becomes *x*_𝒮_*′* _,*i*_and *n*_*i*_ so in general, the two possible maxima for *T*_*l*_(*x*′_𝒮_) are max{*n*_2,*i*_, *x*_𝒮_*′* _,*i*_} and *n*_*i*_.

The same proof holds for *T*_*r*_ (given the symmetry of the expressions), where the maxima is in {max(*x*_𝒮, *i*,_*n*_1,*i*_), *n*_*i*_}.

This proves the lemma.We have reduced the space of possibilities from *O*(*m*^*J*^) to *O*(2^*J*^) (with m the geometric mean of *x*_𝒮_*′*).

We now need to show how to compute this value in *O*(*J* log(*J*)). For this, we rely on the following theorem.

##### Lemma 10

([Papaxanthos et al., 2016]). *Let be a potentially testable subgraph and define* 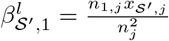 *and* 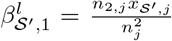, *for j* ∈ {1, …, *J*} *Let π*_*l*_*and π*_*r*_ *be permutations of* 1, …, *J such that* 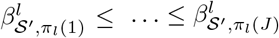*and* 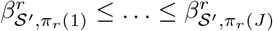, *respectively*.

*Then, there exist κ* ∈ {1, …, *J*} *such that the optimum* 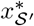 *satisfies either*

- 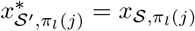 *for j* ≤ *κ and* 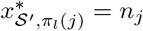 *otherwise* *or*
- 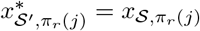 *for j* ≤ *κ and*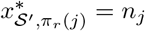 *otherwise*

**Proof of lemma 10** We will do the proof for 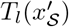. The proof for 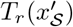 is identical, up to notation.

We can note that since the optimal is at least equal to *n*_2,*j*_, the value of *a*_𝒮,*j*,min_ is *x*_𝒮 *′ j*_ − *n*_2,*j*_. We note 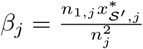. We also note 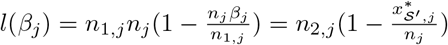. With those notations, we can write

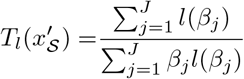

It is straightforward to see that, if 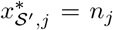, then *l*(*β*_*j*_) = 0. We are then exactly in the setting of Papaxanthos et al. [2016] and we refer the reader to the proof in the supplementary, p.3-6.

This shows that we can compute the envelope in *J* log(*J*)

## S-2 Efficient implementation of CALDERA

We consider a subgraph 𝒮′ created following Algorithm 1, in the second case (Lines 11-15). We therefore have 𝒮, 𝒮_*p*_ such that: 𝒫(𝒮′) = 𝒫(𝒮) = 𝒮_*p*_ and *i*_𝒮_*′* = *i*_𝒮_. After creating 𝒮′, we explore its children, with an itemtable 𝒯. All elements of `Children`(𝒮′, 𝒮_*p*_, *i*_𝒮_*′*, 𝒯) will have a pattern which includes ℐ(𝒮′). Moreover, by definition of the equivalence groups, we already know that {ℐ ∈ 𝒯 : ℐ ⊂ ℐ(𝒮′)} = ∅. Therefore, when constructing 𝒮″ `Children`(𝒮′, 𝒮_*p*_, *i*_𝒮_*′*, 𝒯), only the elements in ℐ(𝒮^″^) \ ℐ (𝒮′) need to be considered.

We store 𝒯 as a matrix of binary patterns. Therefore, some columns can be deleted without loss of information: in Line 13 of algorithm 1, we only keep the columns that are not in ℐ(𝒮). As the `Children` function is called recursively, the itemtable 𝒯 will grow in the number of patterns saved (*i.e* number of rows) but the memory footprint of each pattern will be smaller (*i.e* fewer columns).

## S-3 Background on the minimal p-value

**Figure S1:**
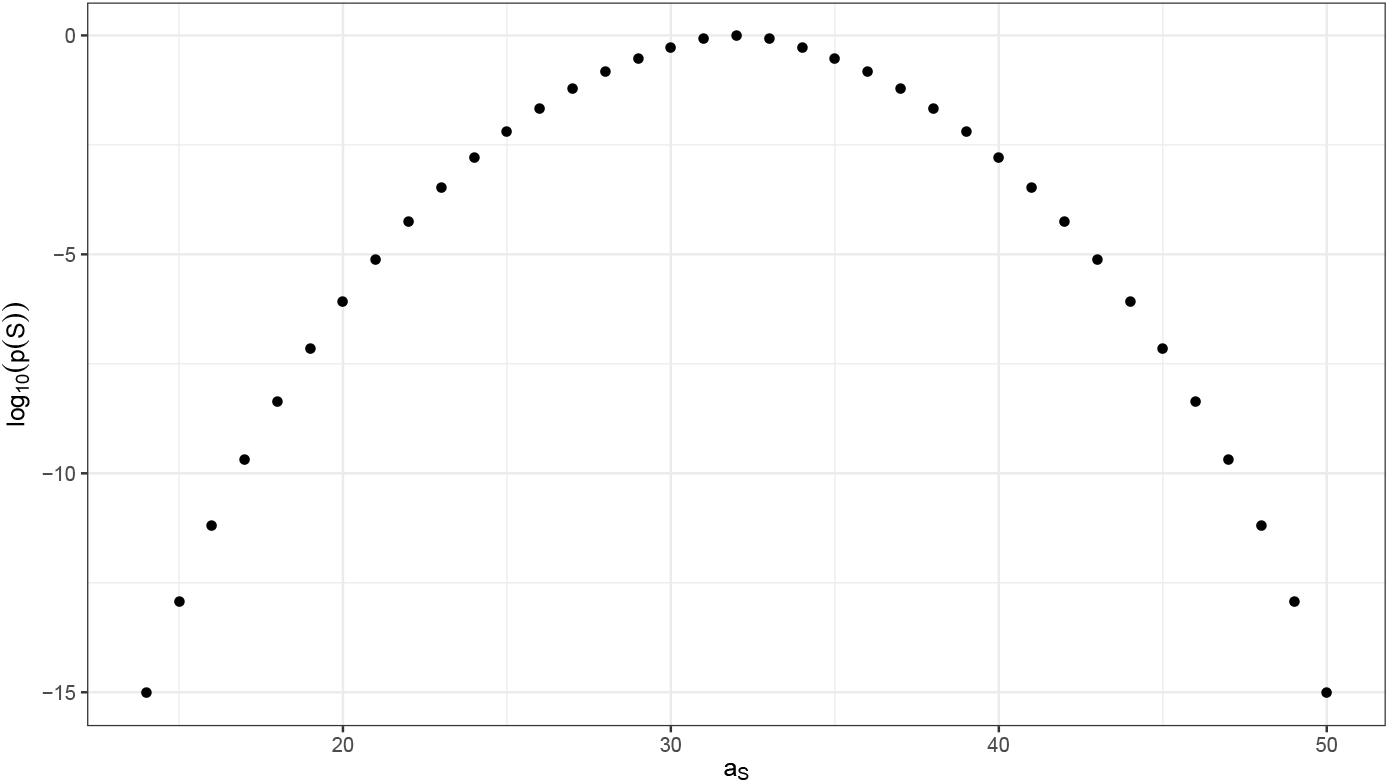
*Finite numbers of possible p-values (log scale) for a fixed value of n*1 = 50 *and x_𝒮_* = 64. *Using the notation from table 1, with J* = 1, *n*_1_ = 50, *n* = 100 and *x_𝒮_* = 64, *the p-value of the χ*^2^ *test is computed for all possible values of a_𝒮_. Since there are only a finite number of possible a_𝒮_ values, there are a finite number of possible p-values, and therefore a smallest one. This minimal p-value can be computed from x_𝒮_, n*1 *and n alone and is* ∼ 10^−15^

## S-4 Benefits of the new envelope

To demonstrate the situations where the new envelope is beneficial, and where it is not, we consider a situation where *n* = 280, *J* = 2. Then, we look at two cases: *n*_1_ = *n*_2_ = 140 and *n*_1_ = 13 × *n*_2_ = 260. In both settings, we compute the envelope as defined in Papaxanthos et al. [2016] and in our case. Then, for *α* = 10^−8^ (a value that we used in practice) and for all possible values of {*x*_𝒮,1_, *x*_𝒮,2_}, we consider whether we would prune (for *k* = 1) using the definition of the envelope from Papaxanthos et al. [2016] or the extended new bound defined in this paper. The new bound nearly doubles the space of prunable subgraphs when there is a clear imbalance, as evidenced in Fig S3b, while it has no effect when the two populations are perfectly balanced, as in Fig S3a.

**Figure S2:**
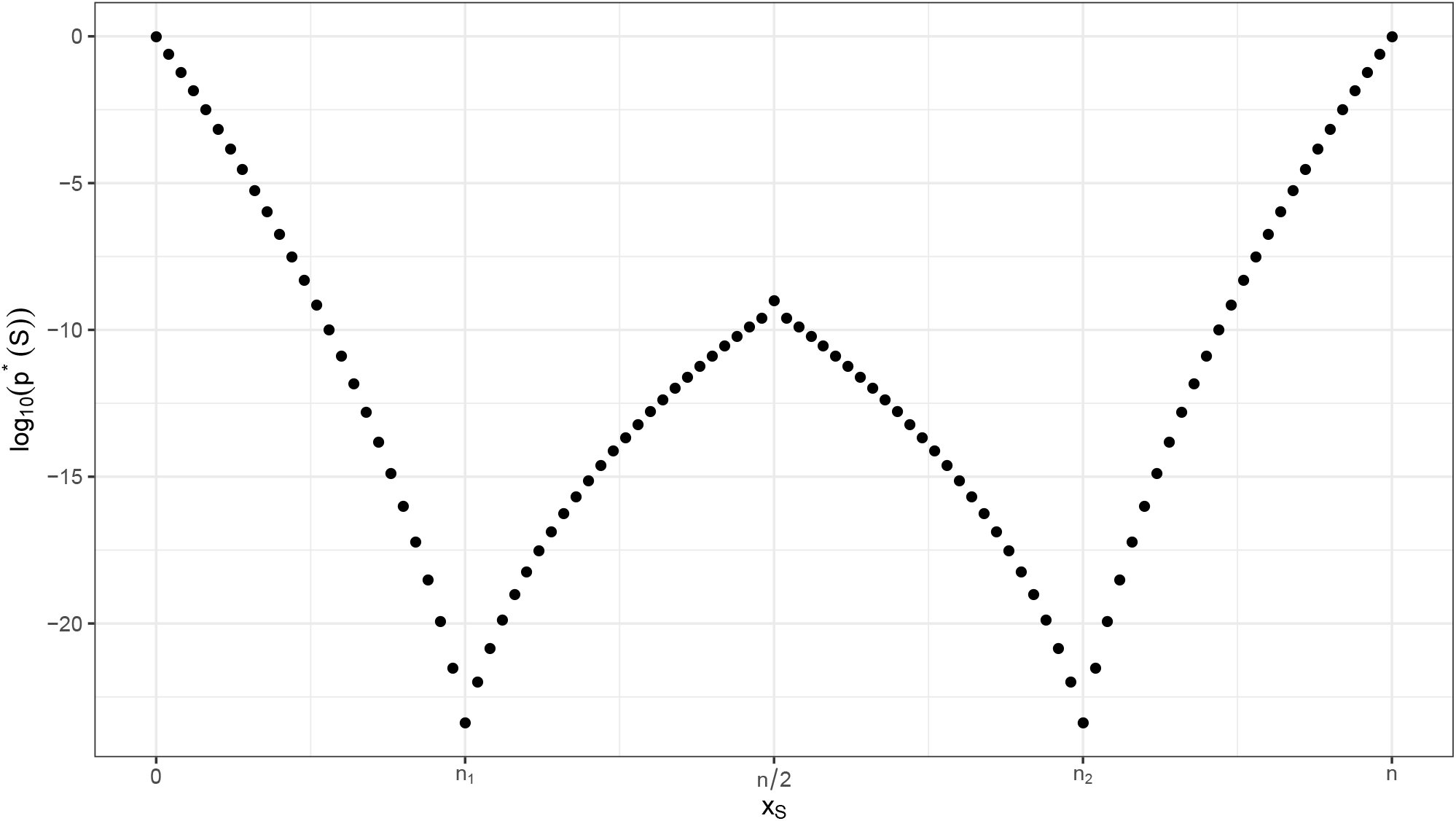
Inspired from Llinares-López et al. [2015] *Minimum p-value as a function of x_𝒮_ for fixed values of n*_1_ = 25 *and n* = 100. *Using the notation from table 1, with J* = 1, *n*_1_ = 25, *n* = 100, *the minimal p-value p^∗^*(*𝒮*) *of the χ*^2^ *test is computed for all possible values of x_𝒮_. For x_𝒮_*≥ max(*n*_1_, *n*_2_), *the minimal p-value is strictly increasing. If we reach that stage, we can prune the graph and stop the exploration in that direction.* *Indeed, if 𝒮*′ ⊇ *𝒮 then x_𝒮_*′ ≥ *x_𝒮_. So if* 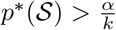, *we know that* 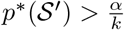 *without computing it*.

## S-5 Benefit of breadth-first search

### S-5.1 A simplified scenario

We consider a very simple graph with *p* = 3 nodes, *J* = 1 population and *n* = 12 samples. The graph is displayed in Fig S4a. Using the reduction from `CALDERA`, we generate a tree structure on 𝒞, displayed in Fig S4b.

Then we can explore this structure in depth-first or breadth-first, while pruning using *α* = 1. The order resulting from an exploration in depth-first can be found in Table S1 and the order fom the exploration in breadth-first can be found in Table S2. In this simple setting, exploring in breadth only visits 4 subgraphs while exploring in depth visits 7. This is because the BFS enumerates testable subgraphs more quickly, thereby increasing *k* and lowering the threshold, which means that the branch starting at {*v*_1_} is pruned earlier in the exploration.

**Table S1:** Order of exploration of the elements of 𝒞 while exploring depth-first

| Subgraph explored | Number of subgraphs explored | Value of the threshold | Testable subgraphs |
| --- | --- | --- | --- |
| $\{v_1\}$ | 1 | .15 | $\{\{v_1\}\}$ |
| $\{v_1, v_2\}$ | 2 | .15/2 | $\{\{v_1\}, \{v_1, v_2\}\}$ |
| $\{v_1, v_2, v_3\}$ | 3 | .15/2 | $\{\{v_1\}, \{v_1, v_2\}\}$ |
| $\{v_1, v_3\}$ | 4 | .15/3 | ∅ |
| $\{v_2\}$ | 5 | .15/3 | $\{\{v_2\}\}$ |
| $\{v_2, v_3\}$ | 6 | .15/3 | $\{\{v_2\}, \{v_2, v_3\}\}$ |
| $\{v_3\}$ | 7 | .15/3 | $\{\{v_2\}, \{v_2, v_3\}, \{v_3\}\}$ |

**Figure S3:**
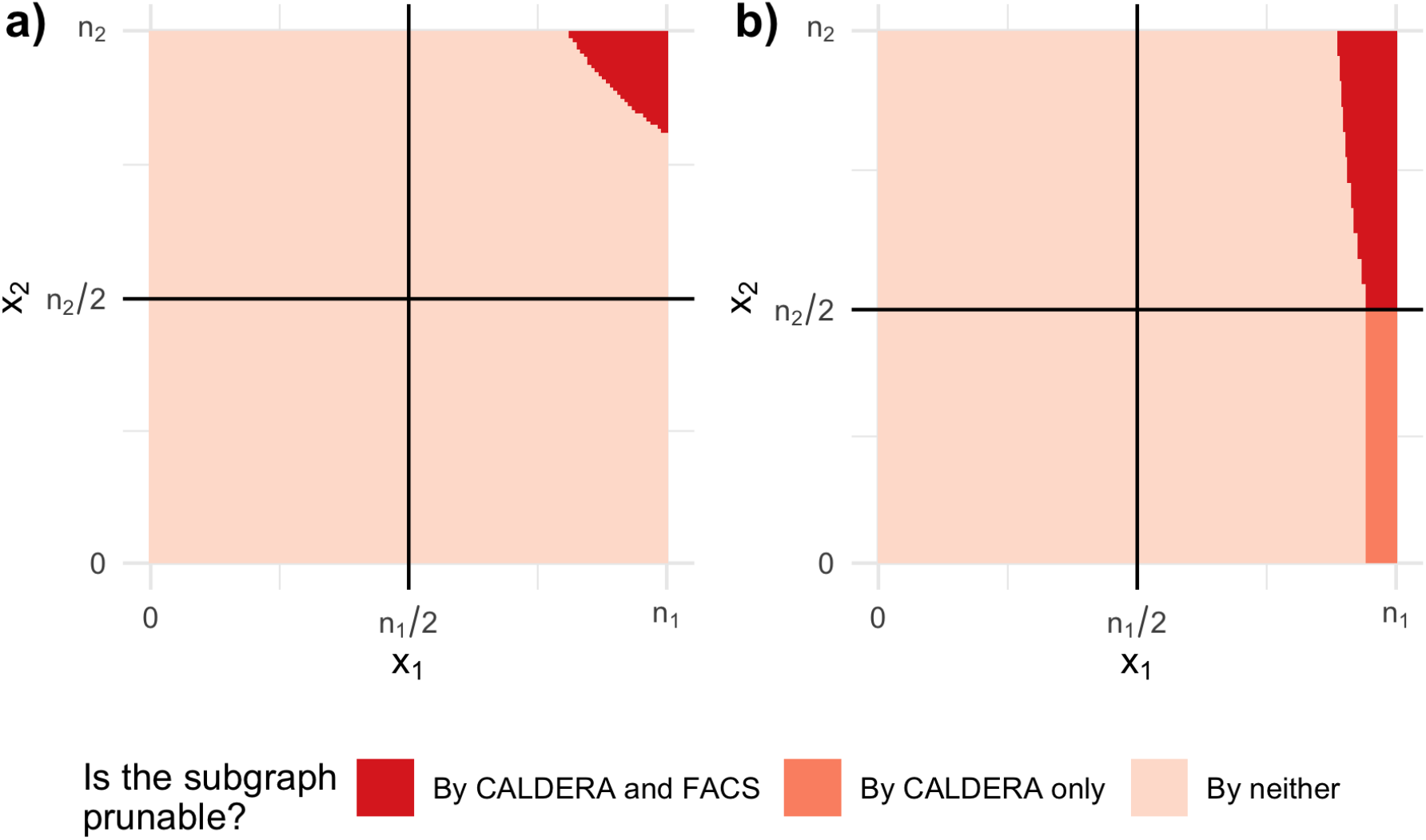
We consider the space of all possible patterns for *n* = 280, *J* = 2 and two cases: a) *n*_1_ = *n*_2_ = 140 and b) *n*_1_ = 260 = 13 × *n*_2_. The phenotypes are well balanced in each population and *alpha* = 10^−8^. The extended lower bound increases the number of prunable subgraphs when the populations are imbalanced.

**Table S2:** Order of exploration of the elements of 𝒞 while exploring breadth-first

| Subgraph explored | Number of subgraphs explored | Value of the threshold | Testable subgraphs |
| --- | --- | --- | --- |
| $\{v_1\}$ | 1 | .15 | $\{\{v_1\}\}$ |
| $\{v_2\}$ | 2 | .15/2 | $\{\{v_1\}, \{v_2\}\}$ |
| $\{v_3\}$ | 3 | .15/3 | $\{\{v_2\}, \{v_3\}\}$ |
| $\{v_2, v_3\}$ | 4 | .15/3 | $\{\{v_2\}, \{v_2, v_3\}, \{v_3\}\}$ |

### S-5.2 A more general setting to understand why BFS is more efficient than DFS

We consider a very simple graph model where, for *v* ∈ V and *i* {1, …, *n*}, *i* ∈ ℐ(*v*) ∼ Binom(prop) and the patterns are independent across nodes. We have no population structure, which means that we consider Fisher’s exact test. For a given level *α*, we want to compute *f* (*α*, prop) = ℙ(*p*⋆({*v*}) *> α*, ∀*v* ∈ V), that is the probability that no subgraph is testable at the first stage of our tree on 𝒞.

Since we consider Fisher’s exact test, there is a bijection between *p*⋆({*v*}) and *x*_{*v*}_ so *p*⋆({*v*}) *> α* ⇒ *x*_{*v*}_ ≥ *σ*(*α*). Moreover, *x*_{*v*}_ ∼ B(prop, *n*), so *f* (*α*, prop) = 1 – (**F**_B_⟩_\_≀⇕_(prop,*n*)_ (*σ*_*α*_))^*p*^with **F**_B_⟩_\_≀⇕_(prop,*n*)_ the cumulative distribution function of the binomial (prop, *n*). Since the nodes are independent, the distribution of *x*_𝒮_ at any stage of the tree can be computed by recursion. We furthermore assume that the graph structure is such that the number of closed subgraphs is *s* × *p* at stage *s*.

In Fig S4c, we display the probability that any subgraph is 1-testable or prunable at stage *s*, for *s* ∈ {1, 2, 3}, *p* = 100 and *α* = 10^−4^.

For most of the range of values, there is at least one testable subgraph in the first stage. So, by exploring in a BFS manner, we start the second stage with a much lower threshold (i.e., a much higher value of *k*) which leads to more pruning. For very low values of `prop`, there might be no testable subgraphs at the first stage but there will be at the second stage, which still justifies an exploration in depth. Note that for large *p*, we can see that there is no testable subgraph at the stages 2 and 3. That is because all such subgraphs have a pattern that is too large. While there may be not testable subgraphs, there are many prunable ones. In that case, an exploration in breadth-first or depth-first would be identical.

This example simplifies two aspects which have opposite effects. The first is that, in practice, the probability of *i* ∈ ℐ(*v*) is of course not uniform across the graph. It is a distribution with much heavier tails which means that, even if the average number of 1 might be small, it is still quite likely that at least one subgraph is testable. The second is that the patterns of neighbouring nodes are correlated. As such, the patterns cannot increase by as much between stages, which limits both the increase in testable pattern discovery, and the pruning.

**Figure S4:**
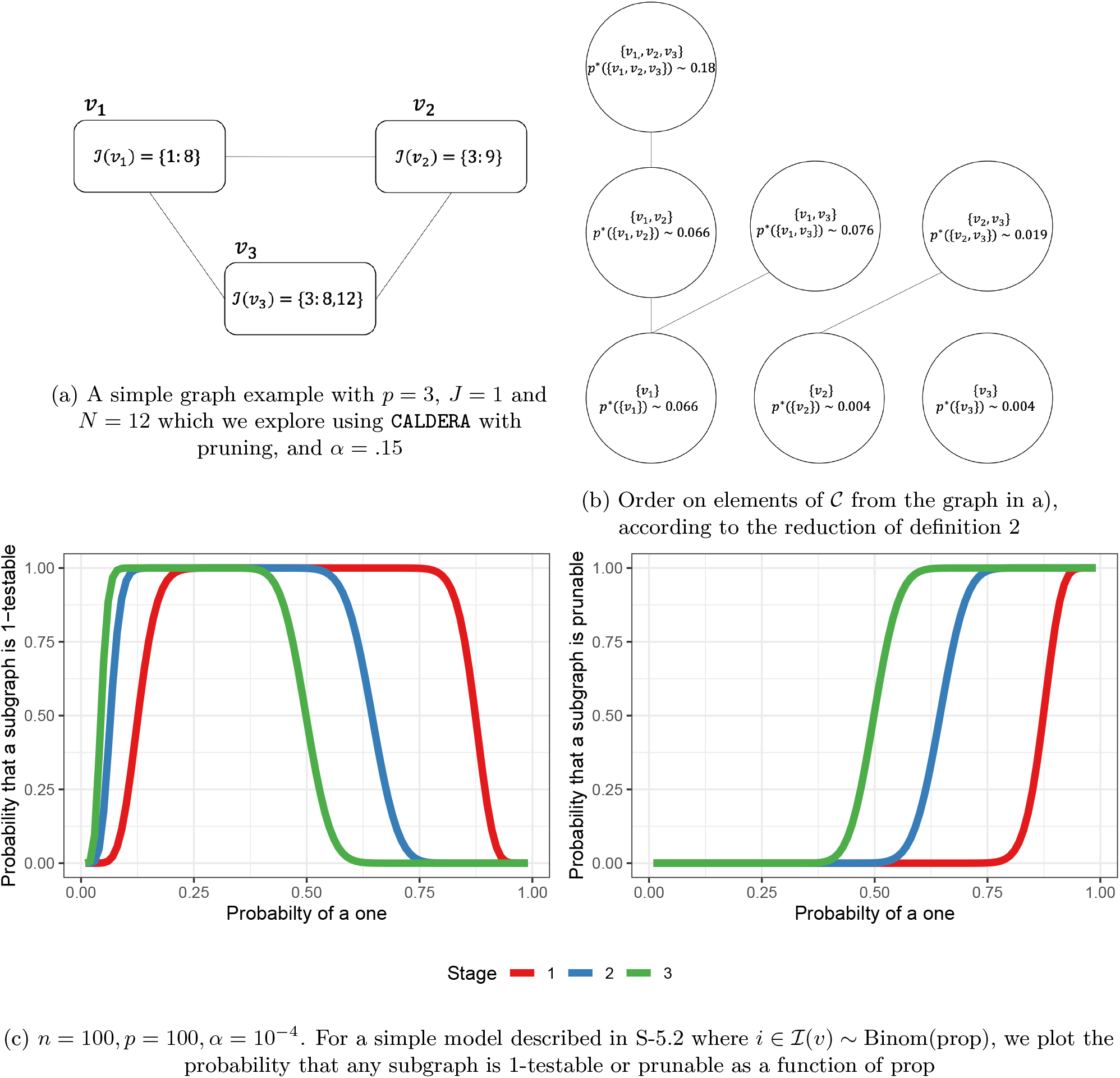
Simple examples where the search in breadth-first is much more efficient that depth-first

## S-6 Speed Simulations

### S-6.1 General simulation settings

For given values of *n* (number of samples) and *p* (number of nodes), we first generate *n* samples with phenotype *y*_*i*_ ∈{0, 1} such that **P**(*y*_*i*_ = 0) = *prop* (user defined parameter). Then, we generate *p* nodes. 10% of the nodes will be associated with the phenotype. For each node in the remaining 90%, we randomly generate 3 edges between this node and another in the 90%. The average degree is therefore 6. For those nodes *v*_*j*_, the associated pattern ℐ (*v*_*j*_) is a random vector such that **P**(*i* ∈ℐ (*v*_*j*_)) = 0.5.

Then, we generate the remaining 10% of the nodes associated with the phenotype. We first generate associated patterns ℐ_*sig*_ such that **P**(*i* ∈ℐ_*sig*_|*y*_*i*_ = 1) = 0.95 and **P**(*i* ∈ℐ_*sig*_|*y*_*i*_ = 0) = 0u.05. Then, those patterns are split into 10 significant nodes sig_*j*_ such that **P**(*i* ∈ℐ (sig_*j*_)|*i* ∈ℐ_*sig*_) = 0.9 and ℐ (∪_j∈[1…10]_)= ℐ_*sig*_

### S-6.2 Parameters for various scenarios

We increase *p* until `COIN+LAMP2` times out (sometimes we went a little further to continue investigation the behaviors).

**Table S3:** Parameter values for the simulations

| Scenario | 1 | 2 | 3 | 4 |
| --- | --- | --- | --- | --- |
| $n$ | 100 | 50 | 100 | 100 |
| $\text{prop}$ | .5 | .5 | .2 | .2 |
| $\alpha$ | .05 | .05 | .05 | $10^{-4}$ |
| timeout (days) | 2 | 1 | 1 | 1 |
| max value of $p$ | $2 \times 10^4$ | $2 \times 10^4$ | $2 \times 10^3$ | $5 \times 10^4$ |

### S-6.3 Results on all scenarios

All computations were run on a r3.4xlarge AWS machine with 16 vCPUs (8 physical ones) and 122GiB[AWS, 2020]. We stop every method once it runs for more than timeout. We also stopped running the entire scenario once we have reached timeout for `COIN+LAMP2` (expect for scenario 3 where we continued further to study the behavior of the various modes of `CALDERA.`

### S-6.4 Memory requirements

We also launched scenario 2 while monitoring memory usage for `COIN+LAMP2, CALDERA BFS` and CALDERA DFS. CALDERA DFS uses 1*/*3 of the peak memory of `CALDERA BFS.` This is expected since the tree structure that is explored scales in *p* in breadth but in *n << p* in depth. Memory-wise, `CALDERA` BFS is on par with `COIN+LAMP2`, which relies on a `DFS` search. This shows that the use of local itemset tables offers memory gains that are enough to offset the exploration in breath, while providing large speed gains. This also suggests that hybrid explorations might be even better at navigating the memory-speed trade-off.

### S-6.5 Imbalance

**Figure S5:**
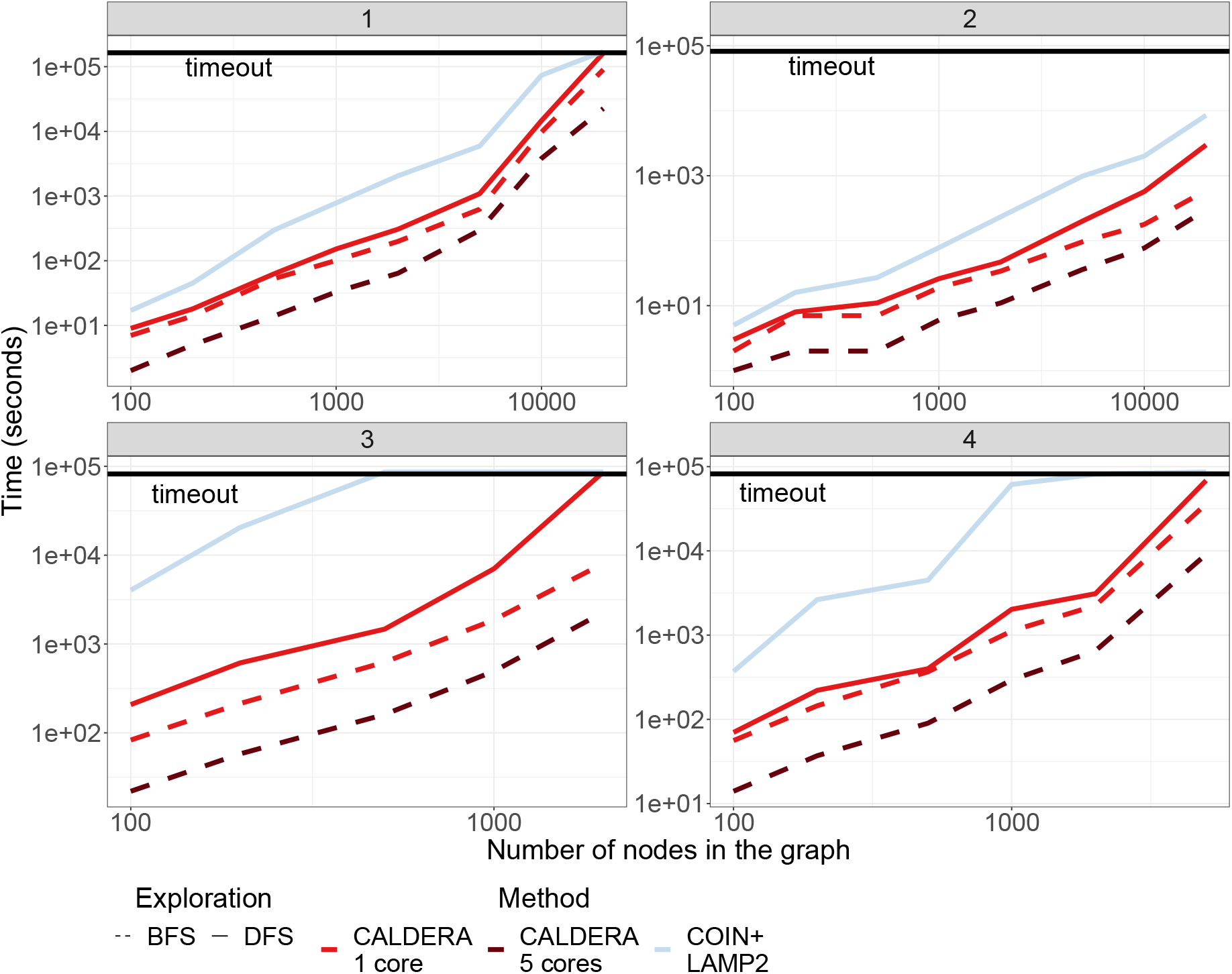
Runtimes for `CALDERA` and `COIN+LAMP` on graphs with various values of covariates *p* and various values of the simulation parameters.

**Figure S6:**
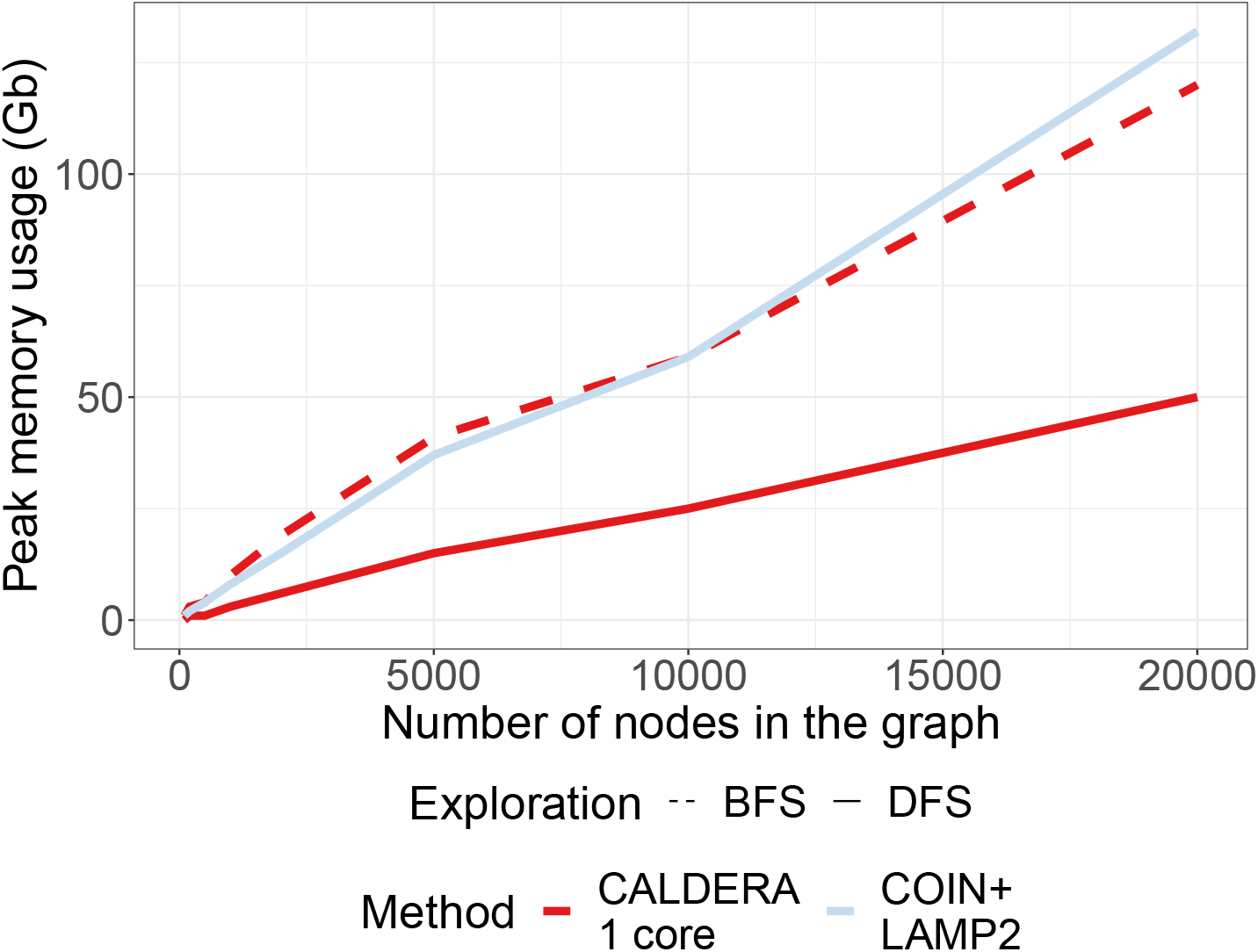
Peak memory usage for `CALDERA` and`COIN+LAMP` on graphs with various values of covariates *p*.

**Figure S7:**
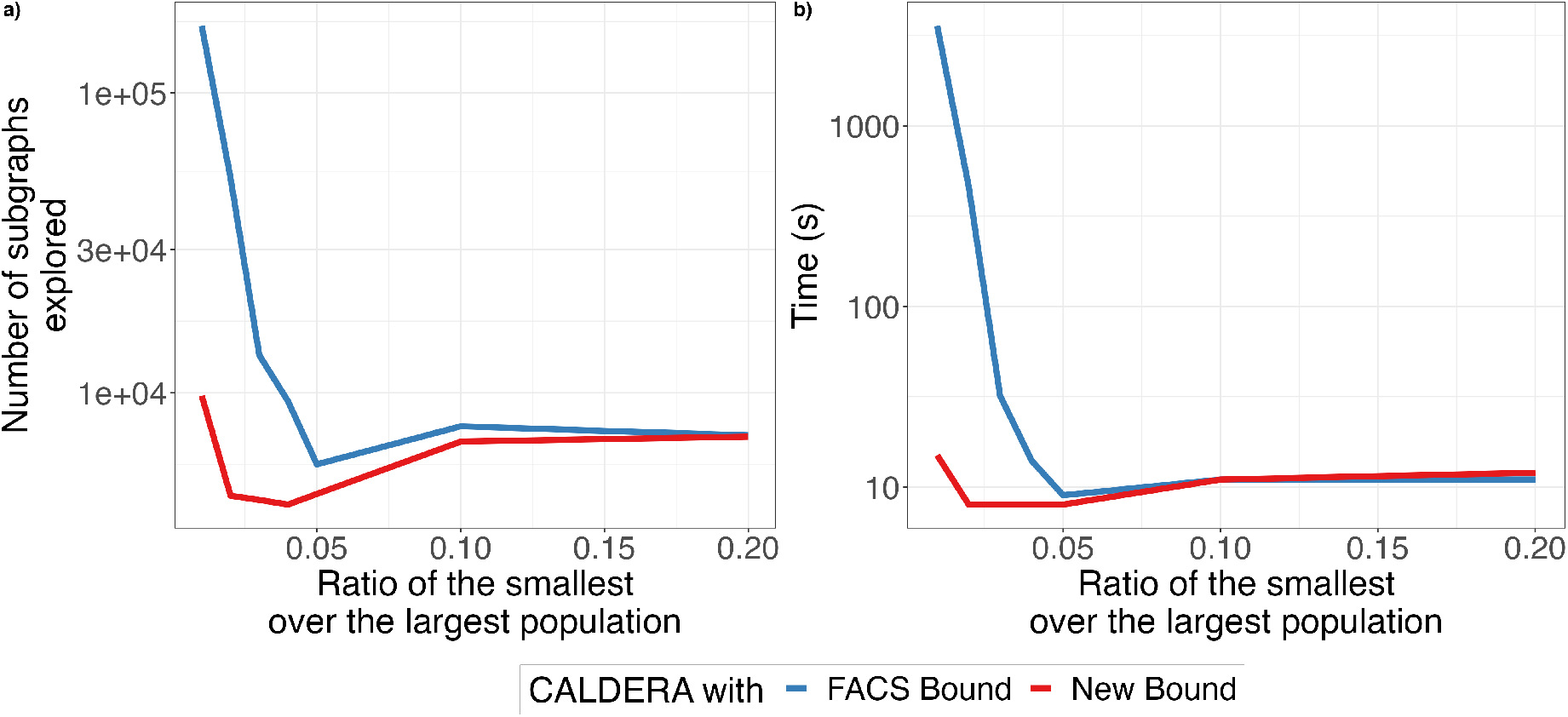
Impact of the new envelope bound on the number of explored subgraphs **a)** and subsequently on runtime **b)** when there is clear imbalance between the size of the two populations.

## S-7 Power Simulations

### S-7.1 Data generation

We first generate two sequences of nucleotides of 1000 bps, gene A and gene B. Then, for each gene, we generate 10 versions. For each version, we start from the original copy and introduce mutations in the following manners: at each base, there is a 1% chance of a point mutation (with each mutation equally likely), a 1% chance of a deletion, a 1% chance of a insertion after the base (with each insertion being equally likely) and therefore a 97% chance that nothing happens. We then generate 50 sensitive samples by randomly selecting with replacement a version of gene B: this is their entire genome. We also generate 50 resistant samples by randomly selecting with replacement a version of gene A and a version of gene B and collating them together. We therefore obtain 100 genetic sequences, 50 of each phenotype.

### S-7.2 Testing all methods

We build a De Bruijn Graph of 704 unitigs using the first step of `DBGWAS`. Then, we run `DBGWAS` with the parameter *-SFF q1.0* to avoid filtering any unitig and then keep those whose p-values are smaller than *α/*704. We also tests the pattern of each unitig for association with the vector of phenotypes. We use the `LAMP2` procedure to remove non-testable hypotheses, at levels of *α*. Finally, we run `CALDERA` with options *–Lmax=2000 -C 1*, and we either specify the maximum number of stages (among [1, 2, 3, 5, 10, 15, 20]) or we run all stages. We look at the list of significant CCSs and keep all their unitigs as significant.

### S-7.3 Results

We compute two metrics: the proportion of unitigs called significant divided by all the unitigs that originated from gene A, named **coverage**; and the proportion of unitigs called significant divided by all the unitigs that originated from gene B, named **False Positive Rate**. We can plot those values for all three methods, and over all ranges of the parameter *–stage* for `CALDERA`

**Figure S8:**
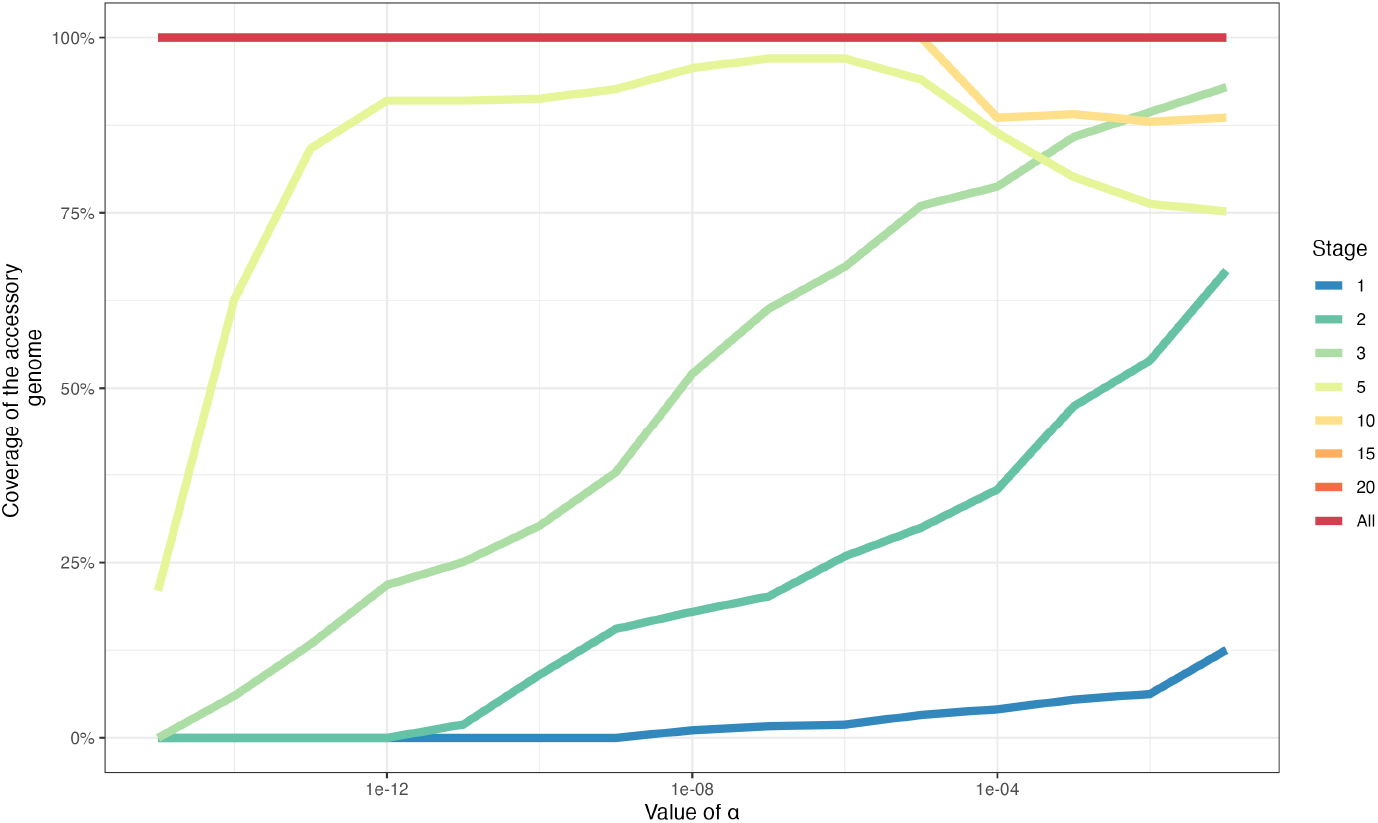
Proportion of all unitigs associated with the resistant phenotype that are found to be significant by `CALDERA` when varying the maximum number of stages explored.

## S-8 Network-guided GWAS on *A. thaliana* genomes

We obtained over 6 millions SNPs and a “date to flowering” phenotype for *n* = 936 *A. thaliana* genomes from easyGWAS [Grimm et al., 2017]. We also obtained 137 *A. thaliana* metabolic pathways from KEGG [Kanehisa and Goto, 2000] using the KEGGrest [Tenenbaum, 2020] and DEGraph [Jacob et al., 2012] R packages. The union of the pathways involved *p* = 3150 genes and the average degree of the resulting graph is ∼ 21.2. We mapped each SNP to the closest gene using snpEff [Cingolani et al., 2012] and defined each gene to be mutated in a sample if it contained at least one mutation mapping to the gene. Runtime was under a minute using 8 cores. *k*_0_ = 70 and 10 subgraphs are found to be significant. In particular, the first and second most significant subgraphs involve pathways ATH00260:*Glycine, serine and threonine metabolism* and 03013: *RNA transport*, which were previously linked to flower development [Hesse and Hoefgen, 2003, Pfaff et al., 2018].

**Figure S9:**
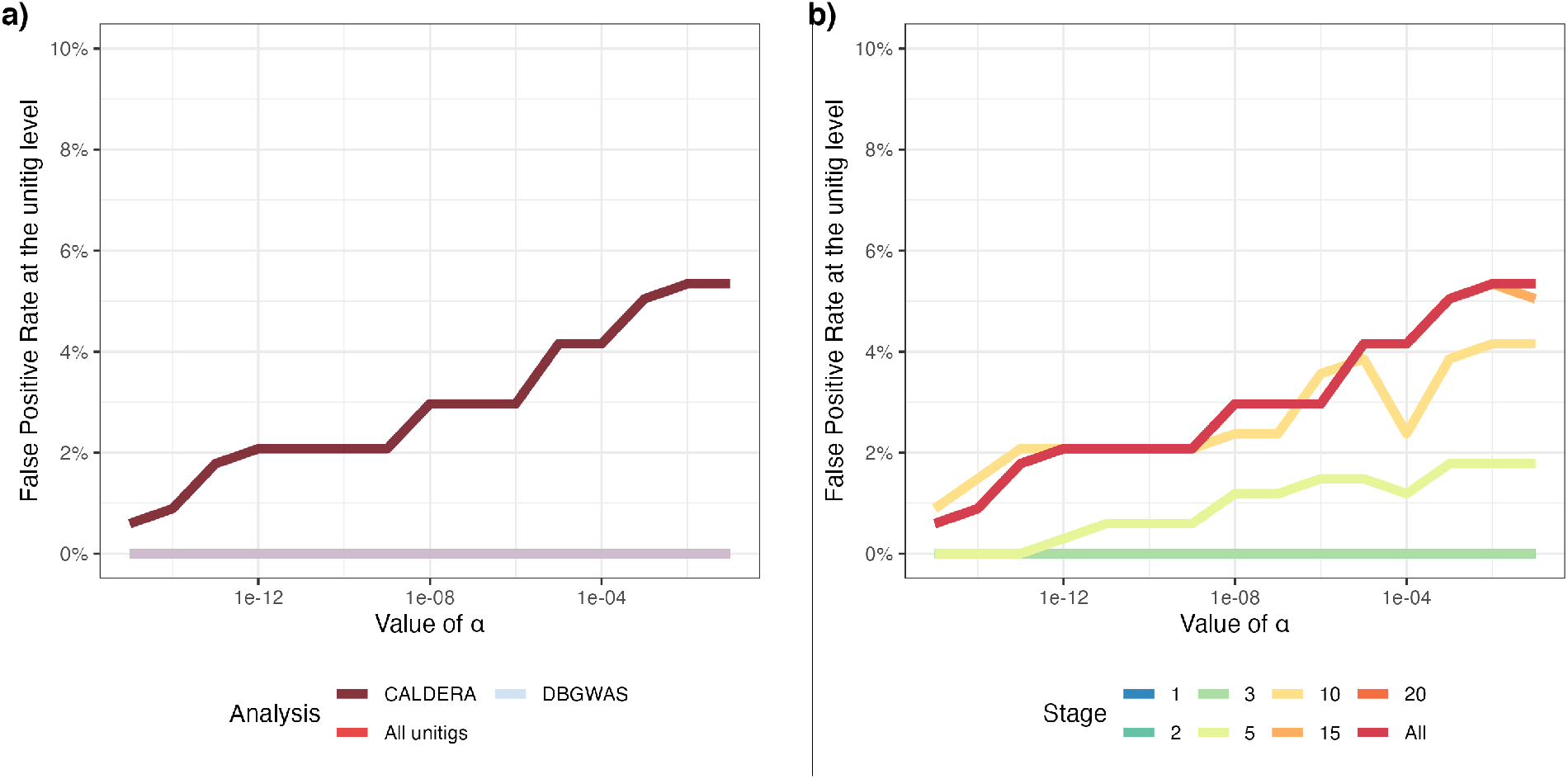
Proportion of all unitigs not associated with the resistant phenotype that are found to be significant as the value of *α* changes **a)** by `CALDERA`, the `LAMP2` procedure on all unitigs and `DBGWAS`, and **b)** by `CALDERA` when varying the maximum number of stages explored.

## S-9 Behavior of CALDERA against BFS stages

Figure S10 shows the number of unitigs belonging to at least one significant CCS for increasing BFS stages on the **Pseudomonas** dataset with *α* = 10^−6^. This number increases sharply at stage 4, where the large CCS containing the plasmid are found, then increases more slowly and reaches a plateau after stage 6.

Figure S11 shows the computational ressources used by `CALDERA` against the BFS stage. Memory usage is exponential in the BFS stage, which is consistent with the number *k*_0_ of explored CCS (not shown), but as seen above, this does not reflect the final coverage of the compacted DBG by significant CCS. The elapsed time increases linearly, with some variation that can be explained by external factors impacting the computation nodes. We do not report the results for stage 8 on the same plot as they were done with a smaller batch size0020of 100, 000 and are therefore not comparable: the corresponding run used 947Gb of RAM and took 44h30.

**Figure S10:**
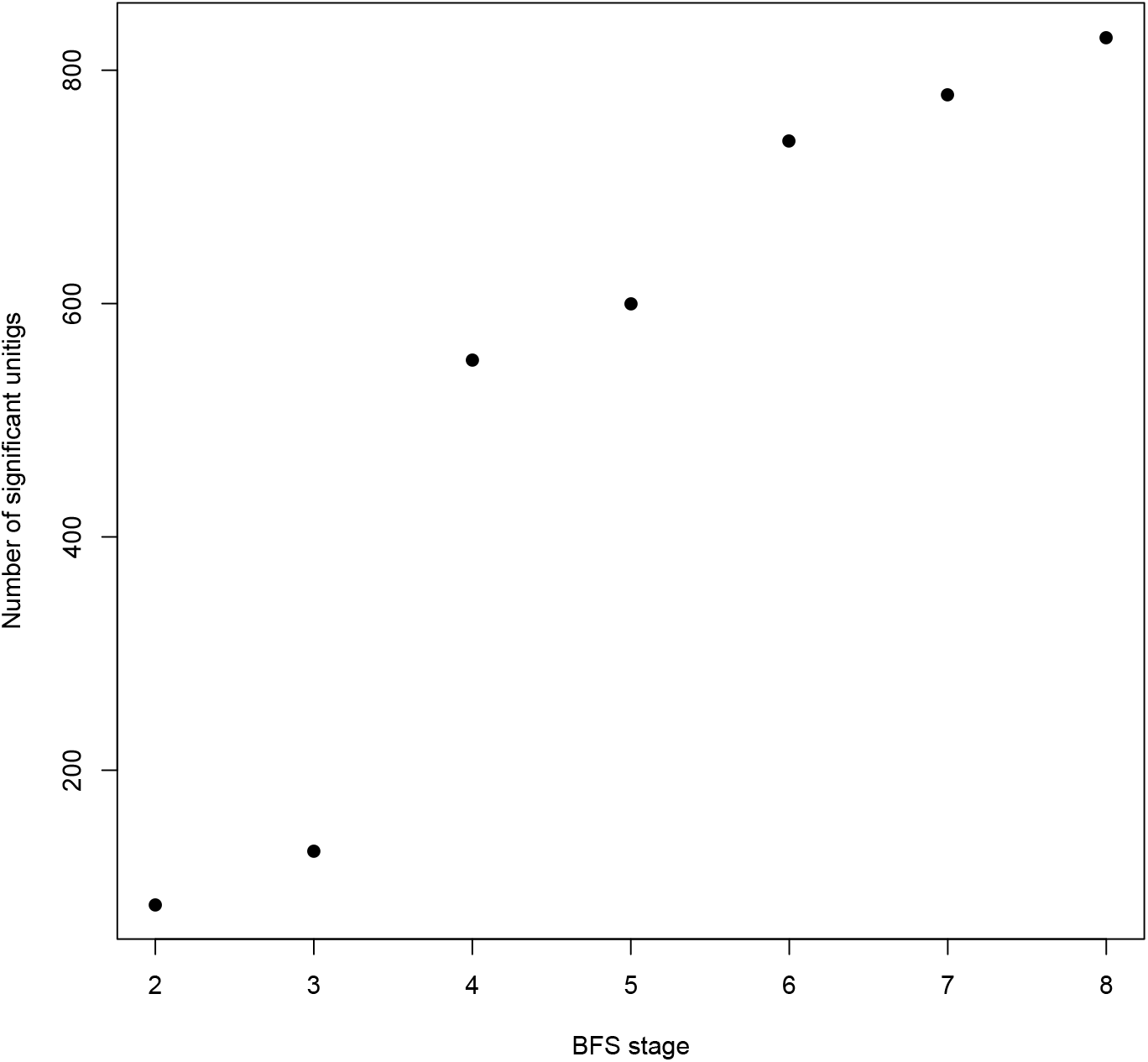
Number of unitigs belonging to at least one significant CCS for increasing BFS stages on the **Pseudomonas** dataset with *α* = 10^−6^.

**Figure S11:**
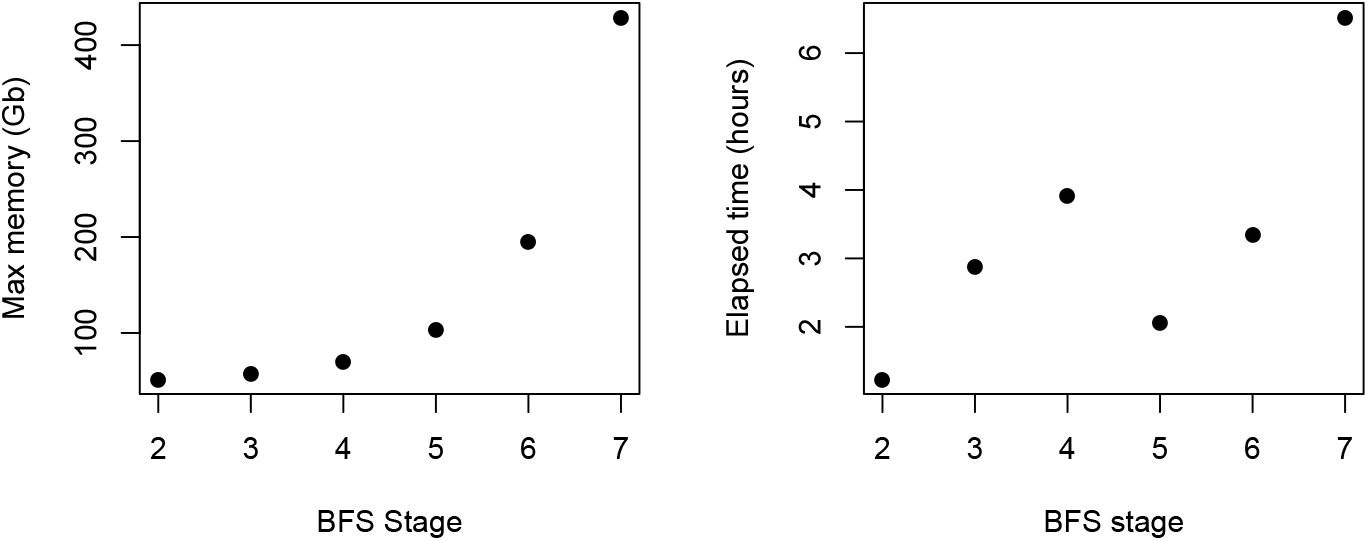
Computational ressources used for increasing BFS stages on the **Pseudomonas** dataset with *α* = 10^−6^, using a batch size of 200, 000 and four cores. Left panel: peak memory used during computation in Gb. Right panel: elapsed time in hours.

